# IP6K1 interacts with the syndecan SDC4 to support secretory granule biogenesis in gastric chief cells

**DOI:** 10.1101/2025.09.17.676719

**Authors:** Jayraj Sen, Pranjali Pore, Rashna Bhandari

## Abstract

Inositol hexakisphosphate kinases (IP6Ks) catalyse the synthesis of the inositol pyrophosphate 5-InsP_7_, and regulate diverse physiological processes. Mice lacking IP6K1 display reduced body weight despite normal food intake, a phenotype that is more apparent in juvenile mice during their rapid growth phase. Additionally, *Ip6k1*^-/-^ mice exhibit decreased serum albumin, elevated faecal protein, and reduced skeletal muscle mass compared to *Ip6k1*^+/+^ mice, suggestive of a deficiency in protein digestion in the absence of IP6K1. We found that IP6K1 is expressed throughout the mouse gastrointestinal tract, and is especially enriched in the cytoplasm of chief cells in the stomach, which are responsible for the storage and secretion of digestive enzymes. Pepsinogen C (PGC) containing granules were sparse, and gastric lipase F (LIPF) granules were completely absent in the gastric glands of *Ip6k1*^-/-^ mice, despite normal expression levels of these enzymes, implicating IP6K1 in digestive enzyme granule biogenesis. Consequently, the level of the active protease pepsin C was decreased in the gastric lumen of *Ip6k1*^-/-^ mice compared with their wild type counterparts. CRISPR/Cas9-mediated deletion of IP6K1 in the gastric adenocarcinoma cell line AGS was able to recapitulate the phenotype of reduced PGC granule intensity seen in gastric chief cells of *Ip6k1*^-/-^ mice. PGC granule formation was restored in *IP6K1^-/-^* AGS cells by the reintroduction of catalytically active or inactive IP6K1, indicating that IP6K1 supports the formation of secretory granules independent of its ability to synthesise 5-InsP_7_. The proteoglycan SDC4, identified as an interactor of IP6K1, was seen to co-localise and co-migrate with PGC granules in *IP6K1^+/+^* but not in *IP6K1^-/-^* AGS cells. Our findings identify IP6K1 as a novel regulator of secretory granule biogenesis in gastric chief cells, to influence protein digestion in the mammalian stomach.

## INTRODUCTION

Inositol hexakisphosphate kinases (IP6Ks) are a family of enzymes that catalyse the synthesis of the inositol pyrophosphate 5-diphosphoinositol pentakisphosphate (5-PP-InsP_5_ or 5-InsP_7_) from inositol hexakisphosphate (InsP_6_) and ATP (1, 2). The three mammalian IP6K paralogues, IP6K1, IP6K2, and IP6K3, differ significantly in their expression patterns and physiological functions (2, 3). IP6K1 and IP6K2 are broadly expressed throughout all mammalian organ systems, whereas IP6K3 expression is predominantly confined to the brain and muscle (1, 4). Engineered deletion of the *Ip6k1* gene (*Ip6k1^-/-^*) in mice and cell line models has revealed its involvement in several cellular and physiological processes (5). Cells depleted for IP6K1 show impaired DNA repair (6, 7), defects in vesicle trafficking (8–11), impaired cell migration (12, 13), altered histone methylation (14), and reduced formation of processing bodies (15), among other changes. In mice, the first study reporting a global deletion of *Ip6k1* described three stark phenotypes - male infertility, reduced serum insulin, and decreased body weight (16). Subsequent studies described the molecular underpinnings of male infertility in these mice, attributing it to cytoskeleton abnormalities in testis architecture and a defect in spermatid maturation (17–19). Extensive work by several groups also revealed different aspects of disrupted insulin homeostasis in *Ip6k1* knockout mice, including impaired insulin secretion in pancreatic β-cells and altered insulin signalling in adipocytes, liver and muscle (20–25). Additional work using the *Ip6k1* knockout mouse model revealed other functions of IP6K1, including roles in blood clotting (26), defence against bacterial pathogens (27, 28), cognition (29, 30), and neuronal migration (13). However, the mechanistic basis of the original observation that *Ip6k1^-/-^*mice maintained on a normal diet have low body weight compared with *Ip6k1^+/+^* mice, has still not been addressed.

As IP6K1 expression has been reported throughout the mammalian gastrointestinal tract (31–33), we wondered if the influence of IP6K1 on body weight may be related to a possible role for this protein in digestion physiology. Digestion of proteins and lipids is initiated in the stomach, with the gastric epithelium playing a crucial role in the release of digestive enzymes. The gastric epithelium comprises a heterogeneous population of surface mucous cells, parietal cells, gastric chief cells, and enteroendocrine cells, which actively secrete protective mucus, hydrochloric acid (HCl), pepsinogen, gastric lipases, and regulatory hormones (34, 35). Gastric chief cells, the principal producers of gastric digestive enzymes including pepsinogens such as pepsinogen C (PGC) and gastric lipase (LIPF), appear basophilic on H&E staining due to their ER-rich cytoplasm (36). Parietal cells, which appear eosinophilic, generate HCl during digestion, which denatures proteins, and also enables the conversion of pepsinogen to pepsin, initiating the breakdown of proteins in the chyme into smaller polypeptides (37). LIPF, which is responsible for 10-30% of lipid digestion, catalyses the breakdown of triglycerides into diacylglycerol (DAG), monoacylglycerol (MAG), and free fatty acids (38, 39). In gastric chief cells, the cellular processes involved in the formation, sorting, and trafficking of secretory granules, such as those containing PGC and LIPF, is understudied (40, 41).

Here, we present evidence that IP6K1, which localises to chief cells in the mouse gastric epithelium, plays a critical role in the formation and secretion of PGC and LIPF containing secretory granules. We observe that in the absence of IP6K1, the expression of PGC and LIPF is unaltered, but their packaging into granules, and stimulated secretion is impaired. Mice deficient for IP6K1 show impaired protein digestion, evidenced by increased retention of protein in faecal pellets from *Ip6k1*^-/-^ mice compared with their *Ip6k1*^+/+^ littermates. Using the gastric adenocarcinoma AGS cell line as a model system, we identified Syndecan-4 (SDC4) (42), a transmembrane proteoglycan, as a specific interacting partner of IP6K1, which acts alongside IP6K1 to facilitate PGC granule assembly and secretion. Our data establishes that IP6K1 acts independently of its catalytic activity to support gastric enzyme secretion, providing the first evidence for the involvement of IP6K1 in mammalian digestion physiology.

## RESULTS

### IP6K1 deletion disrupts body weight homeostasis in mice

It has previously been reported that the loss of IP6K1 leads to a 15-20% reduction in body weight when mice are fed a normal diet (16). This difference is exacerbated as mice age, or are shifted to a high-fat diet (22, 23, 25). These prior studies examined mice between the ages of 4 weeks and 22 months. To determine if the difference in body weight is apparent in juvenile mice, we started monitoring their weight at 16 days postpartum (dpp). Pre-weaned *Ip6k1^-/-^* mice aged between 16 to 22 dpp showed a 15-20% decrease in body weight compared with *Ip6k1^+/+^* mice (Fig. 1A). This difference was more pronounced during the weaning period when mice transitioned from mother-dependent nursing to a conventional diet, with a maximum weight difference of ∼35% seen in 30 ddp mice. In adult *Ip6k1^-/-^* mice (age 45-90 dpp), we continued to observe a marginal yet consistent decrease in body weight and size compared with age-matched *Ip6k1^+/+^* mice (Fig. 1A, B; S1A). As shown earlier (16, 22), we confirmed that there is no significant difference in food intake between *Ip6k1^-/-^* and *Ip6k1^+/+^* mice (Fig. 1C). The decreased weight of *Ip6k1^-/-^* mice has been primarily attributed to their significantly lower levels of body fat compared with *Ip6k1^+/+^* mice (22, 23, 25, 43). These previous studies also monitored serum lipid metabolites in both sets of mice, revealing lower levels of serum triglycerides in *Ip6k1^-/-^* mice that are aged or maintained on a high fat diet (22, 25). We measured serum protein and lipid parameters in a cohort of young adult (∼60 dpp) *Ip6k1^+/+^* and *Ip6k1^-/-^* mice fed on a standard diet, and interestingly, observed a reduction in total serum protein levels in mice lacking IP6K1 (Fig. 1D). We noted no significant differences in any of the other serum parameters tested (Fig. S1 B-K).

**Figure 1:**
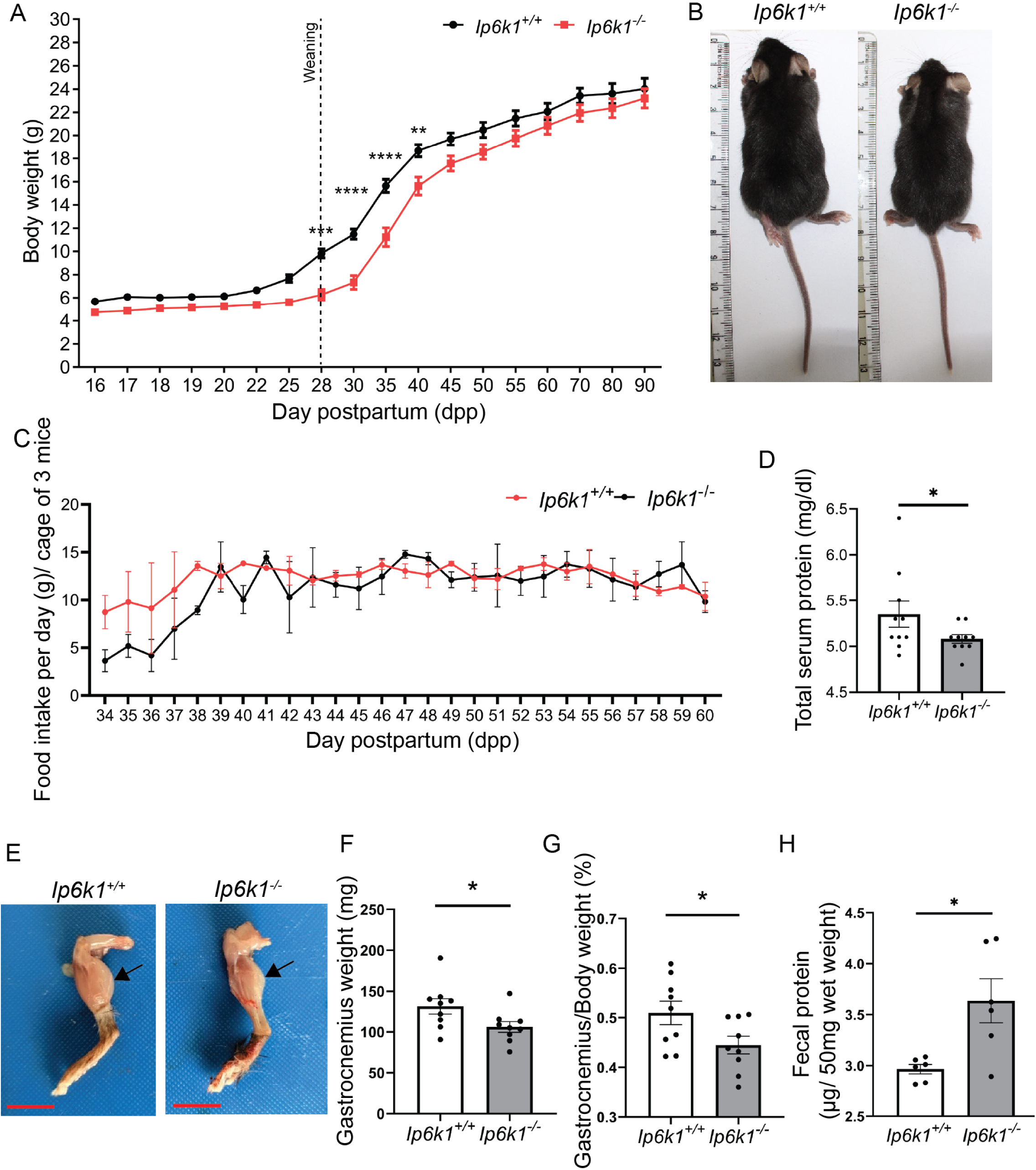
IP6K1 deletion disrupts body weight homeostasis in mice. **A)** Periodic body weight measurement of *Ip6k1^-/-^* and *Ip6k1^+/+^* mice. The body weight was monitored from age 16 days postpartum (dpp) to 90 dpp (Data are mean ± SEM for 16 mice at 16 dpp and 12 mice at 90 dpp for each genotype). Data was analyzed by two-way ANOVA (interaction *P* value is 0.008) with a Bonferroni post-test (*P* value for 28, 30, 35, 40 dpp are 0.0003, <0.0001, <0.0001, 0.0014 respectively). **B)** Representative image of adult mice (male) showing a difference in body size in *Ip6k1^+/+^* and *Ip6k1^-/-^*. **C)** Food consumption in mice to assay the feeding habits in *Ip6k1^-/-^* mice compared with *Ip6k1^+/+^* mice. Data are mean ± SEM (N=6 mice of each genotype). Data was analysed by two-way ANOVA (interaction P value is 0.7742 and is non-significant (ns)). **D)** Total serum protein in adult *Ip6k1^-/-^* and *Ip6k1^+/+^*mice. **E)** Representative images of the isolated lower limb of *Ip6k1^-/-^*and *Ip6k1^+/+^* mice with arrows marking the gastrocnemius muscle. Scale bar 1 cm. **F)** Absolute mass of the gastrocnemius muscle dissected from the lower limbs of *Ip6k1^-/-^* and *Ip6k1^+/+^* mice. Data were analysed using a two-tailed unpaired Student’s *t*-test (mean ± SEM; *N*=10 per genotype, with males and females combined). (\**P* ≤0.05). **G)** Gastrocnemius muscle mass normalized to the body weight for mice in (F). **H)** Faecal protein levels in *Ip6k1^+/+^* and *Ip6k1^-/-^* mice. Data plotted as mean±SEM were analysed using a two-tailed unpaired Student’s t-test (\**P ≤* 0.05; N=6 mice of each genotype).

A decrease in serum protein levels may reflect an overall imbalance in protein digestion or absorption, which could contribute to reduced body weight in *Ip6k1^-/-^* mice. To investigate this possibility, we monitored skeletal muscle mass in these mice, as it is known that protein intake correlates with muscle weight (44, 45). We observed a significant reduction in the weight of the gastrocnemius muscle, even when corrected for reduced body weight, in *Ip6k1^-/-^* mice compared to *Ip6k1^+/+^* mice (Fig. 1E-G). To quantify the extent of protein digestion and absorption, we examined the amount of protein in the faecal pellets from *Ip6k1^-/-^* and *Ip6k1^+/+^* mice housed in metabolic cages (Fig. 1H). We observed significantly higher protein levels in faecal matter from *Ip6k1^-/-^* mice compared to *Ip6k1^+/+^* mice. The increase in undigested faecal protein in *Ip6k1^-/-^* mice points strongly towards impaired protein digestion in these mice.

### IP6K1 is highly expressed in gastric chief cells

In humans and mice, *Ip6k1* mRNA has been detected throughout the gastrointestinal (GI) tract, and tissue immunohistochemistry has confirmed the presence of IP6K1 in glandular cells in these organs (Human Protein Atlas https://www.proteinatlas.org/; Expression Atlas http://www.ebi.ac.uk/gxa)(32, 33, 46). We conducted western blot analysis to confirm the expression of IP6K1 in mouse stomach, duodenum, ileum, colon, and rectum (Fig. 2A). A histopathological examination of these tissues collected from *Ip6k1^+/+^*and *Ip6k1^-/-^* mice did not show any obvious pathophysiological abnormalities (Fig. S2A). There were no significant differences in tissue architecture between the *Ip6k1^-/-^* and *Ip6k1^+/+^* gastrointestinal epithelia, and their cellular composition appeared healthy and normal. A closer look at the gastric glands in the stomach (schematic in Fig. S2B) revealed that the number of gastric chief cells (zymogenic cells), located at the base of the gastric glands, was considerably lower in *Ip6k1^-/-^*compared to *Ip6k1^+/+^* sections (Fig. 2B, C). As the secretion of gastric lipase and pepsinogens by chief cells is crucial for the breakdown of lipids and proteins in the stomach lumen, fewer chief cells may impede gastric digestion in *Ip6k1^-/-^* mice. Immunofluorescence analysis of *Ip6k1^+/+^* mouse stomach sections revealed intense staining for IP6K1 at the base of the gastric glands (Fig. S2C). Within the glands, IP6K1 exclusively showed granular staining in the cytoplasm of chief cells but no staining in parietal cells (Fig. 2D). The high expression levels of IP6K1 in gastric chief cells, coupled with the observed decrease in the number of chief cells in gastric glands of *Ip6k1^-/-^* mice, suggests a role for IP6K1 in mammalian digestion physiology.

**Figure 2:**
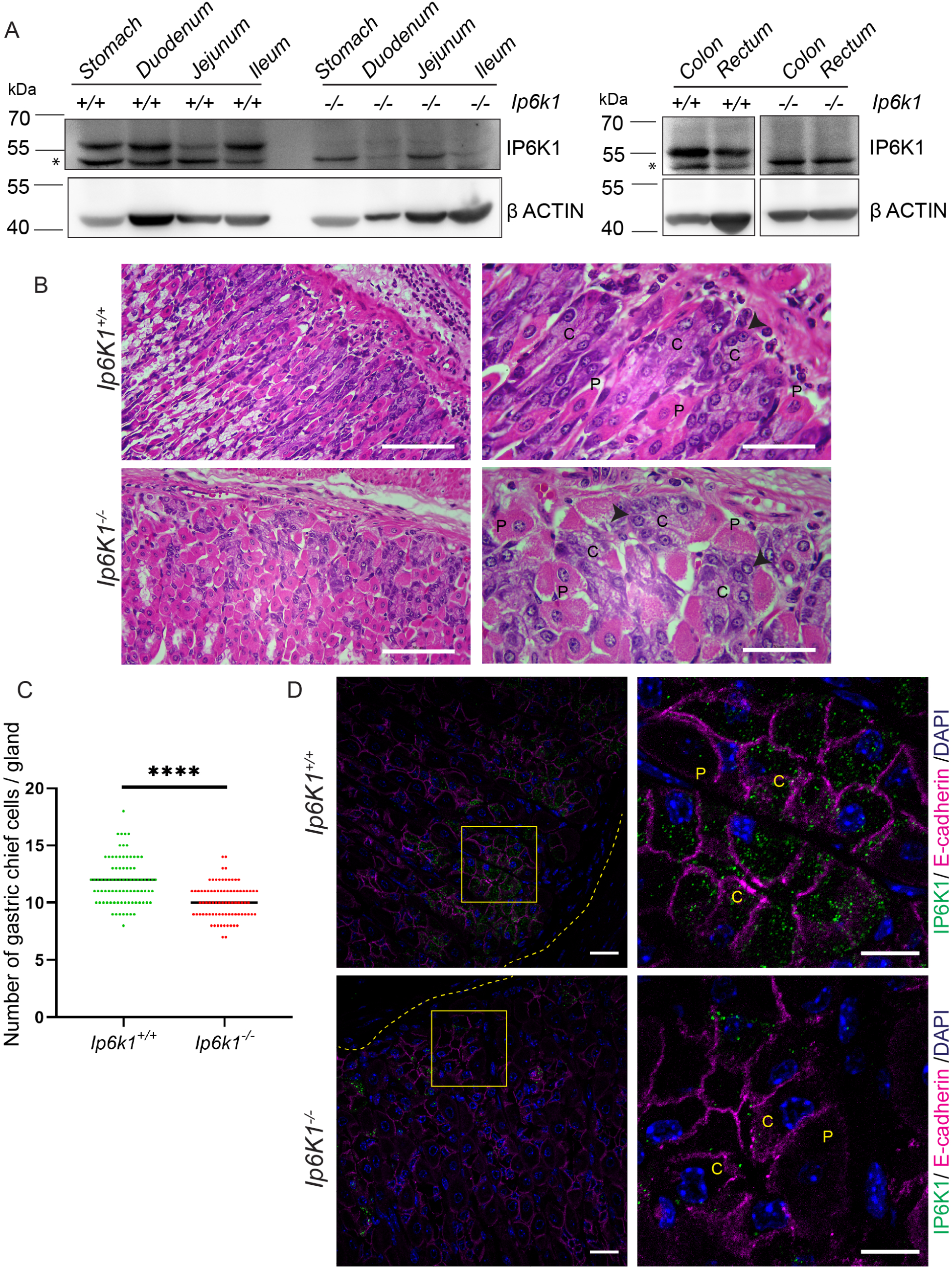
Deletion of IP6K1 in the stomach leads to a reduction in gastric chief cells. **A)** Immunoblot representing IP6K1 expression in adult gastrointestinal tissues (stomach, duodenum, jejunum, ileum, colon, rectum) from *Ip6k1^+/+^* and *Ip6k1^−/−^*mice. β ACTIN was used as a loading control. Expression data presented is representative of N=2 mice for each genotype. The asterisk (*) indicates a non-specific band in these tissues. **B)** Gastric mucosal histology analysis was used to identify different cell types inside the stomach of *Ip6k1^+/+^* and *Ip6k1*^-/-^mice. Haematoxylin stains nuclei and eosin stains the cytoplasmic region of cells (images are representative of N=3 mice of each genotype). Scale bars: left 100 μm, and right 50 μm. Chief cells and parietal cells are indicated by “**C**” and “**P**” respectively in H&E-stained sections. **C)** Quantification of stomach chief cells in the gastric glands of *Ip6k1^+/+^* and *Ip6k1^−/−^* mice. Scatter plots with mean value represent data from 86 glands in *Ip6k1^+/+^* mice and 82 glands in *Ip6k1^−/−^* mice, assessed from three independent age-matched mice of each genotype. Data were analysed using a two-tailed unpaired Student’s t-test (\*\*\*\**P≤* 0.0001). **D)** Representative images of *Ip6k1^+/+^* and *Ip6k1^-/-^* stomach FFPE sections stained to detect IP6K1 (green) expression in gastric glands. E-cadherin staining (magenta) was used to mark the cell boundaries, and DAPI (blue) to mark nuclei. A yellow box marks the regions in the immunofluorescence images on the left that are zoomed in on the right; gastric chief cells (**C)** and parietal cells (**P**) are indicated. E-cadherin staining is unaltered in *Ip6k1^-/-^* mouse stomach. IP6K1 shows granular cytoplasmic staining in chief cells, with specific staining observed in *Ip6k1^+/+^* mouse tissue sections and some non-specific staining detected in *Ip6k1^-/-^* sections. Images, representative of N=3 mice of each genotype, were captured using a Zeiss LSM700 confocal microscope with 63X/1.4 NA objective; Scale bar 20 µm. All confocal immunofluorescence images are z-stacks showing xy dimensions as a maximum intensity projection (MIP).

### IP6K1 deletion impairs accumulation of digestive enzymes in chief cell granules

In an earlier study we have shown that *Ip6k1^-/-^* mouse embryonic fibroblasts display defects in Golgi architecture arising from impaired dynein-driven vesicle transport (8). To examine if the loss of IP6K1 affects Golgi morphology in gastric chief cells, we stained stomach sections to detect the cis-Golgi marker GM130. We observed an increase in the distance between the Golgi network and the basal membrane in chief cells in *Ip6k1^-/-^* compared with *Ip6k1^+/+^* gastric glands (Fig. S2D, E), indicative of altered Golgi architecture in the absence of IP6K1. Next, we examined whether the loss of IP6K1 affects the accumulation of digestive enzymes in gastric chief cells. We stained mouse stomach sections to detect the zymogen pepsinogen C (PGC), the major pepsinogen in adult mice. In *Ip6k1^-/-^* mice, we observed ‘hollow’ PGC containing granules within gastric chief cells, present mostly at the apical region of the cell, contrasting with the uniform distribution of intensely stained PGC granules in stomach sections of *Ip6k1^+/+^* mice (Fig. 3A). Quantification of the immunofluorescence images revealed a reduction in the fluorescence intensity of PGC granules in *Ip6k1^-/-^* compared to *Ip6k1^+/+^* gastric glands, reflecting the hollow appearance of granules in *Ip6k1^-/-^* sections (Fig. 3B). Immunofluorescence analysis of stomach sections to detect gastric lipase F (LIPF) revealed failure of LIPF accumulation in secretory granules in the stomach of *Ip6k1^-/-^* mice, whereas some chief cells in *Ip6k1^+/+^*glands showed LIPF staining (Fig. 3C). Additionally, we found that the enzyme activity of LIPF in stomach lysates of *Ip6k1^-/-^* mice was significantly lower compared to *Ip6k1^+/+^* mice (Fig. 3D). We conducted immunoblotting analyses of stomach lysates to examine whether reduced granule accumulation of PGC and LIPF was due to a reduction in their expression levels. Interestingly, we observed no significant difference in the levels of PGC or LIPF in stomach lysates from *Ip6k1^+/+^* and *Ip6k1^-/-^* mice (Fig. 3E, F and G), suggesting that these enzymes fail to accumulate in secretory granules despite their normal levels of expression in IP6K1 depleted chief cells. Together, these data suggest that impaired vesicle trafficking arising from the absence of IP6K1 in chief cells, which is manifested as anomalous Golgi morphology, could compromise the accumulation of digestive enzymes in secretory granules despite their normal expression levels.

**Figure 3:**
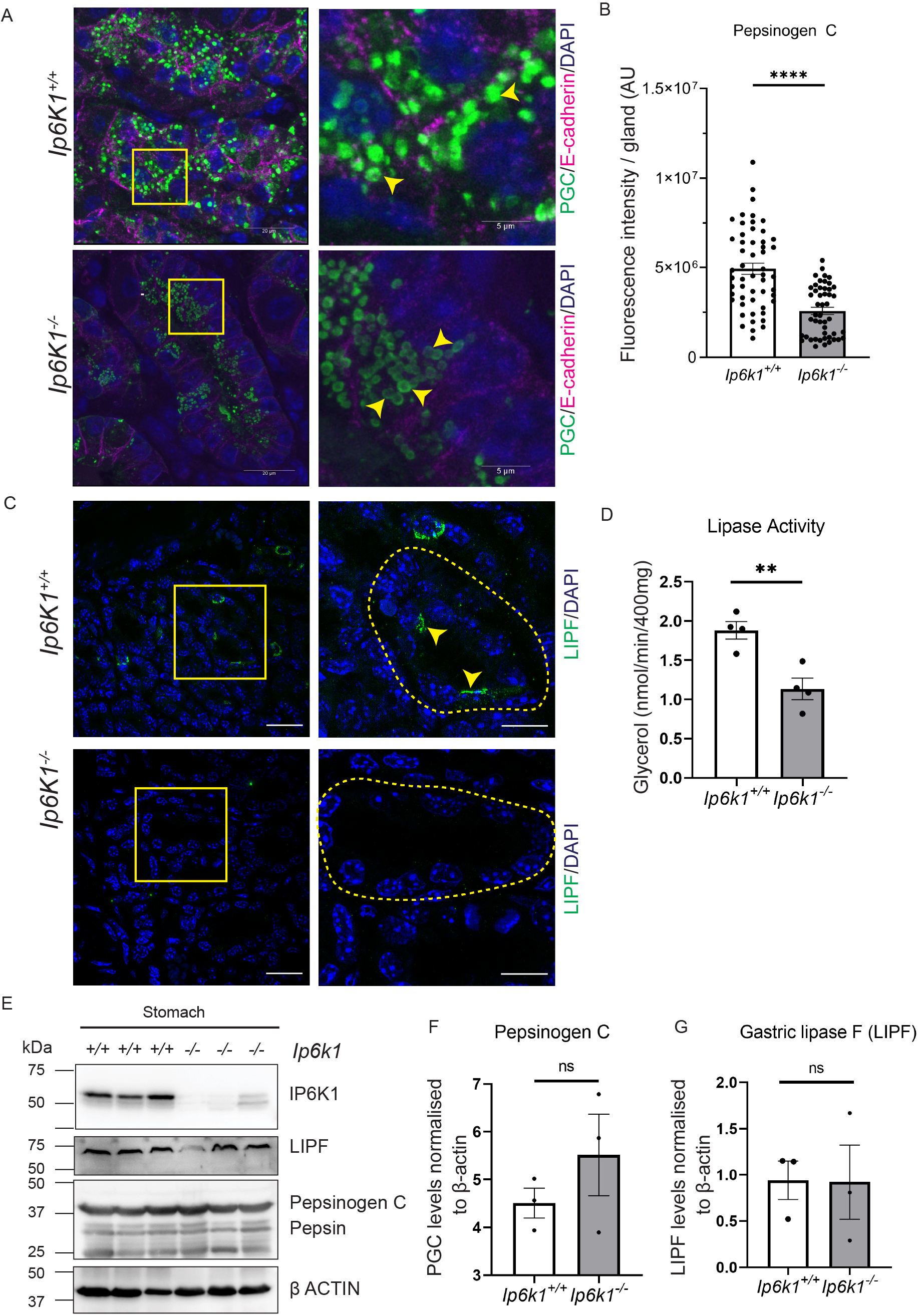
IP6K1 deletion impairs accumulation of digestive enzymes in chief cell granules. **A)** Immunofluorescence staining of stomach FFPE sections from *Ip6k1^-/-^* and *Ip6k1^+/+^* mice to detect PGC (green) containing secretory granules. E-cadherin antibody (magenta) was used to mark the cell boundaries and DAPI (blue) marked the nuclei. A yellow box marks the regions in the immunofluorescence images on the left that are zoomed in on the right. Scale bars: left 20 µm; right 5 µm. Images were captured with a Leica TCS SP8 confocal microscope using a 63x 1.4 NA oil immersion objective. Yellow arrowheads mark PGC containing granules in chief cells. Images are z-stacks showing xy dimensions as a maximum intensity projection (MIP). **B)** The graph represents the quantification of fluorescence intensity of PGC staining in the stomach of *Ip6k1^+/+^* and *Ip6k1^-/-^* FFPE sections. N=53 glands in *Ip6k1^+/+^*and 48 glands *Ip6k1^-/-^* stomach FFPE sections from three mice of each genotype. Data were analysed using a two-tailed unpaired Student’s *t-*test *(**** P≤ 0.0001)*. **C)** Representative immunofluorescence images of gastric epithelium from FFPE sections of *Ip6k1*^+/+^ and *Ip6k1*^-/-^ stomach stained for the chief cell marker LIPF (gastric lipase F) (green) (indicated by yellow arrowheads). Nuclei were stained with DAPI (blue). A yellow box marks the regions in the immunofluorescence images on the left that are zoomed in on the right, and a dashed line marks the boundary of a gastric gland in the image on the right. Scale bars: left 20 µm; right 10 µm. Images, representative of N=3 mice of each genotype, are z-stacks showing xy dimensions as a maximum intensity projection (MIP). Images were captured using an Elyra 7 (SIM) module of the Zeiss LSM 980 confocal microscope with 63X/1.4 NA Objective. **D)** Gastric lipase (LIPF) activity was analysed using a colorimetric assay in stomach lysates from *Ip6k1^+/+^* and *Ip6k1^-/-^* mice. Data (mean±SEM; N=4 mice of each genotype) were analysed by a two-tailed unpaired Student’s t-test (\*\**P≤* 0.005). **E)** Representative immunoblots of the stomach homogenized in RIPA buffer to check the levels of gastric chief cell-specific markers PGC and LIPF in adult *Ip6k1^+/+^* and *Ip6k1^-/-^* mice (N=3 mice of each genotype). β-actin was used as a loading control. **F and G)** Quantification of data in (E). PGC and LIPF levels in *Ip6k1^+/+^* and *Ip6k1^-/-^* mice in whole cell lysate, normalized to the levels of β-actin in the same lysate. Data (mean ±SEM; N=3 mice for each genotype) was analysed using a two-tailed unpaired Student’s t-test (ns P>0.05).

### IP6K1 regulates the luminal secretion of gastric contents in response to stimulation

Next, we probed the impact of IP6K1 depletion on gastric secretion. We deprived mice of food for 14-16 hours and then surgically ligated the pyloric sphincter which connects the stomach to the duodenum (Fig. 4A). After recovery from pyloric ligation, secretagogues were administered to the mice subcutaneously. We used histamine, which stimulates release of HCl from parietal cells, and carbachol, an acetylcholine receptor agonist, which stimulates granule release from chief cells along with acid release from parietal cells. PBS was administered as a control. Mice were euthanized 2 hours after stimulation and the stomach was dissected to measure gastric pH in situ. As expected, there was a significant decrease in the pH of gastric juice in *Ip6k1^+/+^* mice upon stimulation with either histamine or carbachol (Fig. 4B). Notably, neither secretagogue induced a decrease in gastric pH in *Ip6k1^-/-^* mice. Although there was no detectable expression of IP6K1 in parietal cells (Fig. 2D), the systemic loss of IP6K1 in these mice may have an indirect impact on histamine- or carbachol-stimulated signalling that results in the release of H^+^ and Cl^-^ ions. Subsequent to pH measurement in dissected stomachs, the gastric luminal contents were collected and relative levels of pepsin C were quantified by western blotting. We observed a significant reduction in pepsin C levels in the gastric releasate of *Ip6k1^-/-^* mice compared with *Ip6k1^+/+^* mice upon treatment with carbachol (Fig. 4C, D). There was no difference in basal pepsin C levels in the gastric juice of *Ip6k1^+/+^* and *Ip6k1^-/-^* mice administered PBS or histamine, neither of which stimulate the release of pepsinogen from chief cells. Decreased pepsin C levels in the gastric juice of *Ip6k1^-/-^* mice may be attributed to reduced PGC accumulation in secretory granules of *Ip6k1^-/-^* gastric chief cells (Fig. 3A, B), and higher gastric pH in these mice, which in turn reduces the conversion of PGC to pepsin C. This reduction in pepsin C levels in the stomach of *Ip6k1^-/-^* mice may lead to impaired gastric protein digestion, which correlates with our observation of increased faecal protein levels in these mice.

**Figure 4:**
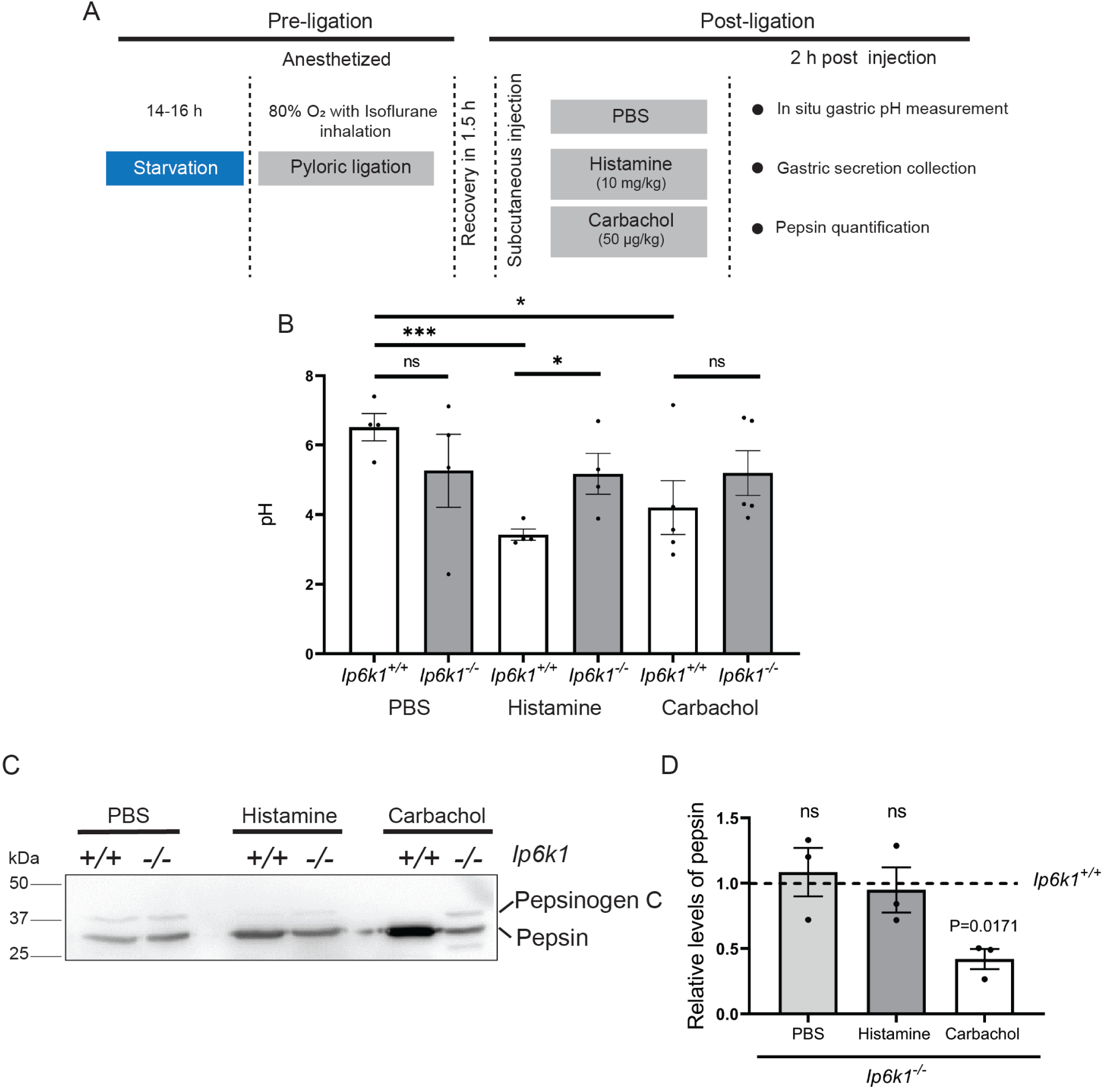
IP6K1 regulates the luminal secretion of gastric contents *in vivo* in response to stimulation. **A)** The schematic describes the experimental workflow for pyloric ligation, timeline for stimulation with secretagogues, and analysis of gastric luminal content. **B)** Graphs show pH of the gastric contents following treatment with PBS, histamine, or carbachol after pyloric ligation, measured using a hand-held pH meter. Data were analysed using a two-tailed unpaired Student’s t-*test (*** P≤0.001; *P≤0.05 and ns P≥ 0.05)*. N=3-4 mice of each genotype. **C)** Immunoblots to detect the levels of pepsin (produced from PGC) in the gastric luminal secretion followed by stimulation with histamine (10 mg/kg body weight), carbachol (50 mg/kg body weight), or PBS as a control. Equal amounts of protein from *Ip6k1^+/+^* and *Ip6k1^-/-^* samples (quantified using BCA assay) were loaded for each treatment. **D)** Quantification of data in (C) showing mean ± SEM with the fold change in levels of pepsin in the gastric lumen in *Ip6k1^-/-^* compared to *Ip6k1^+/+^* mice upon the indicated stimulation. N=3 of three mice of each genotype. Data were analysed using a one-sample t-test (value for *Ip6k1^+/+^* mice treated with PBS, Histamine, and carbachol was set at 1).

### IP6K1 supports PGC granule formation independent of its catalytic activity

To characterize the molecular mechanism by which IP6K1 regulates PGC granule formation, we utilized the human gastric adenocarcinoma cell line AGS, which has been used as a model system to study the formation of secretory vesicles in gastric chief cells. We generated a CRISPR/Cas9-based IP6K1 knockout AGS cell line (*IP6K1*^-/-^), and a control *IP6K1*^+/+^ line was generated using a non-targeting sgRNA construct (Fig. 5A). Immunofluorescence analysis to detect IP6K1 in wild type AGS cells showed the presence of endogenous IP6K1 in peri-nuclear cytoplasmic granule clusters (Fig. 5B). The specificity of this staining was confirmed by its absence in *IP6K1*^-/-^ cells. This granular staining of IP6K1 in AGS cells is in contrast to the predominantly nuclear and nucleolar localisation of IP6K1 observed in HEK293T and U-2 OS cell lines (31, 47), suggesting that IP6K1 localises to different subcellular compartments depending on the cell type and tissue context.

**Figure 5:**
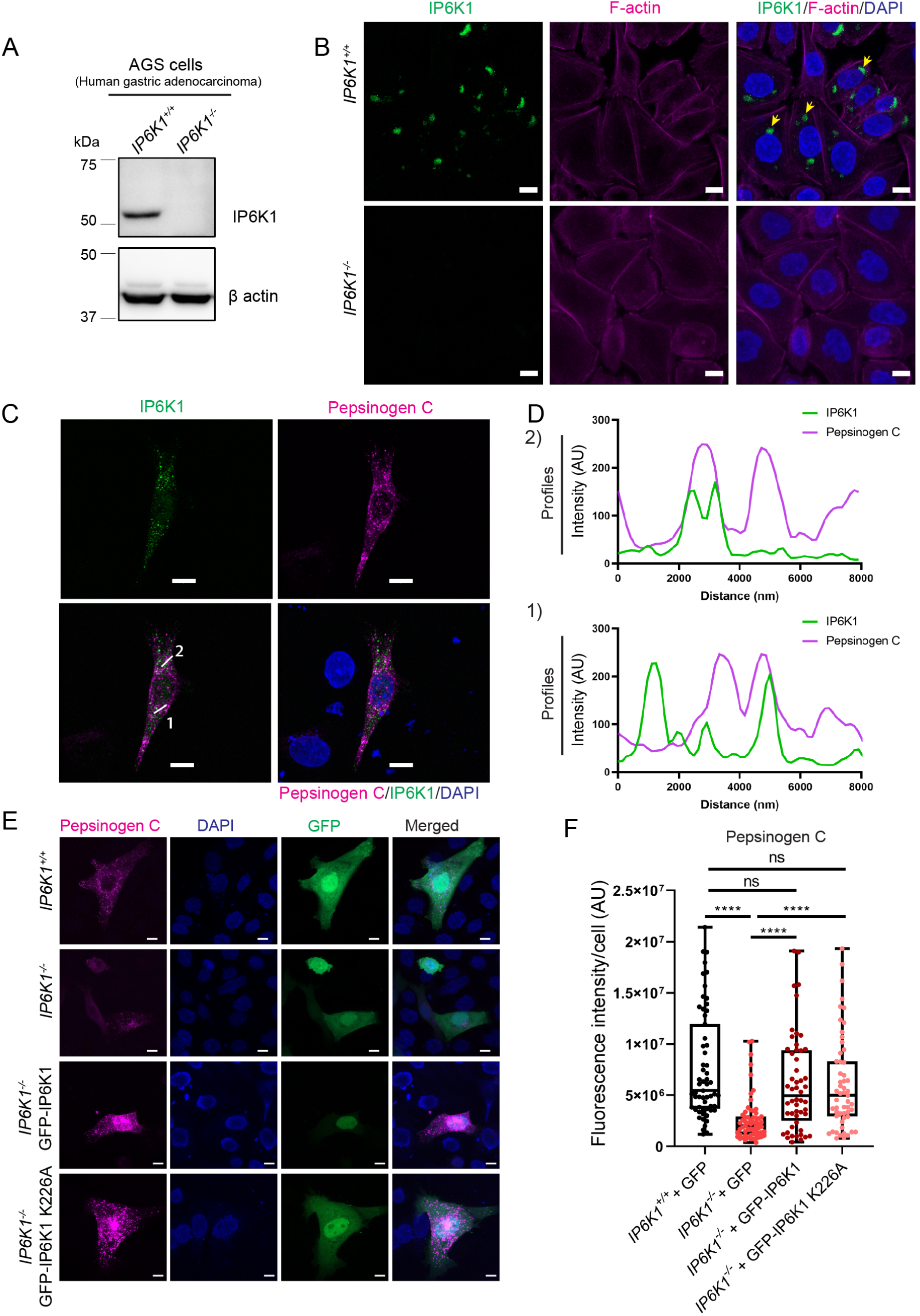
IP6K1 supports PGC granule formation. **A)** Immunoblotting to detect IP6K1 in cell lysates from CRISPR/Cas9 generated single-cell derived *IP6K1^-/-^* and non-targeted control (*IP6K1*^+/+^) AGS lines, using β-actin as a loading control (N=4). **B)** Immunofluorescence analysis to detect endogenous IP6K1 in *IP6K1*^+/+^ and *IP6K1^-/-^* AGS cell lines. IP6K1 (magenta) was present in peri-nuclear cytoplasmic granules in control AGS cells and absent in *IP6K1^-/-^* cells. F-actin was stained using rhodamine-phalloidin to mark the cytoskeleton and identify individual cells, and DAPI (blue) was used to stain the nuclei. Scale bar 20 μm. All confocal immunofluorescence images in (B) are presented as z-stacks with xy dimensions displayed as a maximum intensity projection (MIP). Images were captured with a Leica TCS SP8 confocal microscope using a 63x 1.4 NA oil immersion objective. **C)** AGS wild-type cells co-expressing N-terminally tagged SFB-IP6K1 (green) and C-terminally tagged PGC-mCherry (magenta). Nuclei were stained with DAPI. Images, showing individual channels and an overlay, were captured with a Leica TCS SP8 confocal microscope using a 63x 1.4 NA oil immersion objective; scale bar 10 µm. **D)** Traces show fluorescence intensity profiles for IP6K1 (green) and PGC (magenta), measured along the indicated white lines (1 and 2) in (C), analysed using imageJ. **E)** Control (*IP6K1*^+/+^) or *IP6K1^-/-^* AGS cells overexpressing GFP-IP6K1, GFP-IP6K1 K226A (catalytically inactive), or pEGFP control (green), and co-expressing PGC-mCherry (magenta). Nuclei were stained with DAPI (blue). Scale bar, 10 μm. **(F)** The box-and-whiskers plot shows fluorescence intensity per cell (in arbitrary units, AU) from the rescue experiment. Sample sizes were N=60 and 63 cells for AGS *IP6K1*^+/+^ and *IP6K1^-/-^* cells expressing eGFP and PGC-mCherry, respectively; and N=52 and 55 cells for *IP6K1^-/-^* AGS cells expressing GFP-IP6K1 or GFP-IP6K1(K226A), respectively, along with PGC-mCherry. Data were compiled from three independent experiments. Data were analysed using a Kruskal-Wallis test with Dunn’s post-hoc test (*P* values are indicated; ns non-significant *P* ≥0.05). Confocal immunofluorescence images are z-stacks showing xy, yz and xz dimensions as a maximum intensity projection. All images were taken by a Zeiss LSM700 confocal microscope using a 63X/1.4 NA objective. Images in (E) were subjected to uniform ‘levels’ adjustment in the ZEN software to improve visualization.

To characterise the impact of IP6K1 depletion in AGS cells we first compared Golgi architecture in *IP6K1^+/+^* and *IP6K1^-/-^*AGS cells by immunofluorescence staining to detect GM130. In control AGS cells, the Golgi was seen to be distributed all around the nucleus, whereas in cells lacking IP6K1, there were fewer Golgi clusters in the peri-nuclear region (Fig. S3A, B). This altered Golgi morphology in *IP6K1^-/-^*AGS cells is reminiscent of the defective Golgi architecture observed in *Ip6k1^-/-^*mouse gastric chief cells (Fig. S2D, E). To study the impact of IP6K1 on PGC granule formation, we over-expressed C-terminally mCherry-tagged PGC in AGS cells. PGC localised to cytoplasmic granules in AGS cells, and showed partial co-localisation with granules containing over-expressed IP6K1 (Fig. 5C, D). Next, we examined whether the reduction in PGC granules in *Ip6k1*^-/-^ mouse gastric chief cells (Fig. 3A, B) is recapitulated in the AGS model system. Indeed, AGS cells lacking IP6K1 showed a significant decrease in the fluorescence intensity of PGC-mCherry granules compared with control cells, despite equal levels of expression of the protein (Fig. 5 E, F; S3C). IP6K1 primarily acts by catalysing the synthesis of 5-InsP_7_, but also contributes to cellular pathways independent of its enzyme activity (15, 48). To determine whether 5-InsP_7_ synthesis by IP6K1 is essential to support the formation of PGC granules, IP6K1^-/-^ AGS cells were transfected to express catalytically active or inactive (K226A) forms of EGFP-tagged IP6K1 along with PGC-mCherry (Fig. 5E). For comparison, we expressed EGFP with PGC-mCherry in control or *IP6K1^-/-^* AGS cells. The expression of either active or catalytically inactive IP6K1 was able to restore the formation of PGC granules in IP6K1^-/-^ cells to the level of control AGS cells (Fig. 5F), revealing that IP6K1 works independently of 5-InsP_7_ synthesis to promote the biogenesis of PGC containing secretory granules.

### IP6K1 deletion alters carbachol-induced PGC granule dynamics

Next, we used IP6K1 depleted AGS cells to explore the mechanism by which IP6K1 supports carbachol-stimulated pepsin release in the mouse gastric lumen. We expressed PGC-mCherry in *IP6K1^+/+^* and *IP6K1^-/-^* cells, and treated the cells with carbachol for 30 min to stimulate granule exocytosis. *IP6K1^+/+^* cells stimulated with carbachol exhibited a decrease in their PGC granule content 30 min post-treatment (Fig. 6A, B). Conversely, in *IP6K1^-/-^* cells, where the PGC granule content is low to begin with, there was a significant increase in granule accumulation post carbachol treatment, especially at the periphery of the cells towards the plasma membrane. These observations suggest a possible defect in granule exocytosis in addition to impaired granule formation in IP6K1 depleted AGS cells. To explore this further, we monitored the temporal dynamics of PGC granules for up to 16 min post carbachol stimulation by time-lapse imaging. We plotted kymographs to visualise the movement of individual PGC granules over time (Fig. 6C, D; Video S1, S2). Granule movement in *IP6K1^+/+^* cells was steady and sustained, with higher oscillatory movement reflected in thicker trajectories in the kymograph, whereas in *IP6K1^-/-^* cells, PGC granule dynamics was markedly impaired, displaying reduced mobility and increasing stagnation over time. Quantification of the total distance travelled, and overall displacement of individual PGC granules during the 16 min period of imaging post carbachol treatment revealed a significant decrease in both parameters in *IP6K1^-/-^* compared with *IP6K1^+/+^*cells (Fig. 6E). This reduction in PGC granule mobility may underlie retention of these granules near the cell periphery in *IP6K1^-/-^* cells upon carbachol treatment (Fig. 6A, B).

**Figure 6:**
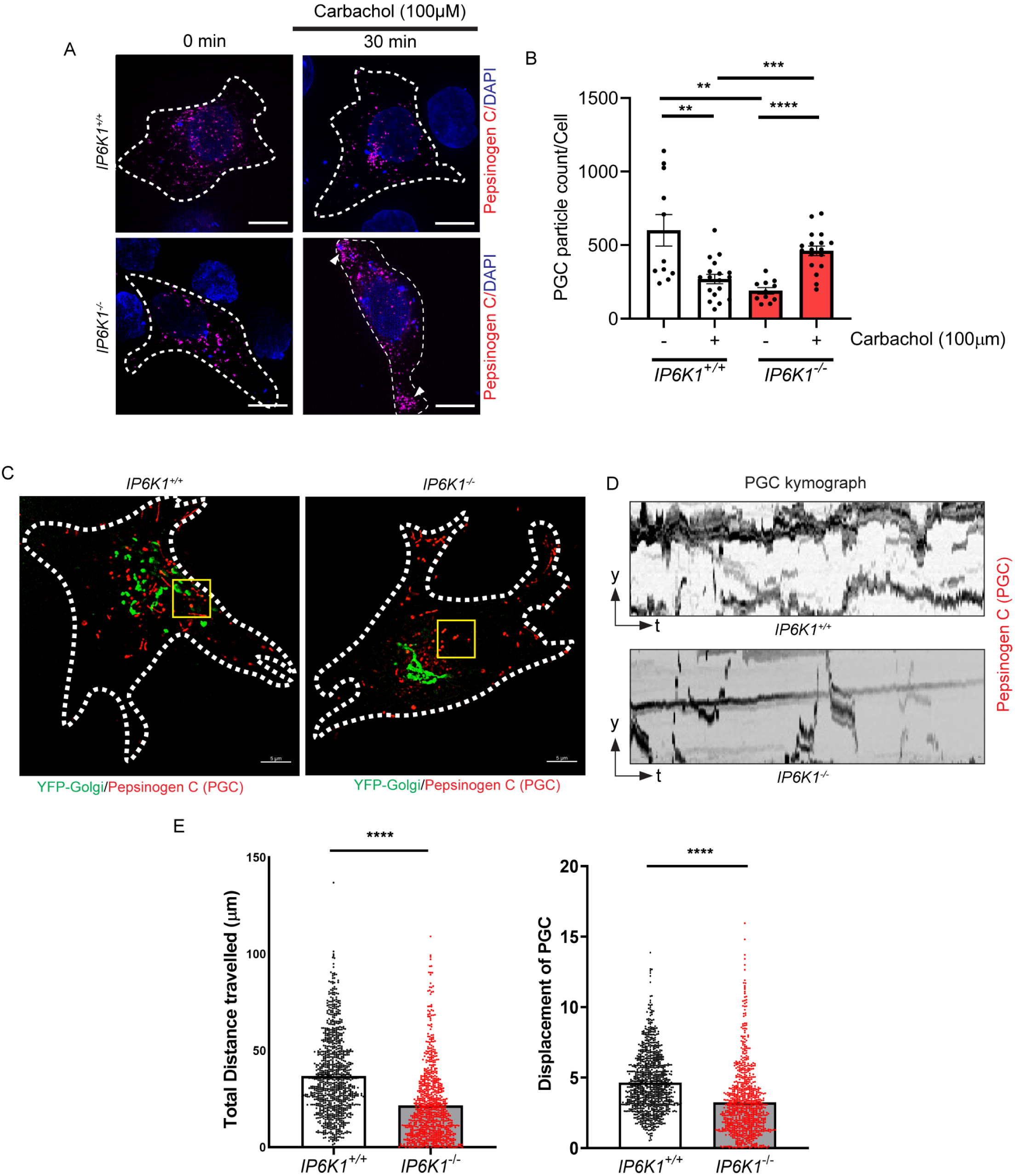
IP6K1 deletion impairs PGC granule formation and dynamics. **A)** Control (*IP6K1^+/+^*) or *IP6K1^-/-^* AGS cells overexpressing PGC-mCherry (magenta) were stimulated with carbachol (100µM), then fixed and imaged at 0 and 30 min. Nuclei were counterstained with DAPI (blue). Scale bar, 10 μm. White arrowheads indicate PGC granule accumulation towards the cell periphery of *IP6K1^-/-^*cells after 30 min of carbachol treatment. **B)** Quantification of the data shown in (A). The graph depicts the number of PGC granules per cell, counted using the particle analysis tool in Fiji. Sample sizes were N=11 cells at 0 min and N=18 cells at 30 min for both *IP6K1*^+/+^ and *IP6K1^-/-^*AGS cells. **C)** Live cell imaging was performed to investigate PGC granule exocytosis in response to carbachol stimulation in *IP6K1^+/+^* and *IP6K1^-/-^* AGS cells expressing YFP-Golgi (green) and PGC-mCherry (red). Cells were stimulated with carbachol (100µM) and imaged at 1/3.8 sec over a period of 16 min (total 200 frames). The representative image was captured at 16 min. Box shows the region in which PGC granules were selected at time 0 min for tracking by kymograph analysis. Scale bar 5µm. Images were captured using the Elyra 7 (SIM) module of the Zeiss LSM 980 confocal microscope with 63X/1.4 NA Objective. **D)** Kymographs (plotted using the Kymograph Builder tool in Fiji) show a comparison of the movement of PGC granules in *IP6K1^-/-^*and *IP6K1^+/+^* cells in response to carbachol stimulation. The y-axis shows the granule location coordinates and the x-axis represents time. Each kymograph shows granule oscillation and movement during the 16 min imaging period. Regions of *IP6K1^+/+^* and *IP6K1^-/-^* are expanded on the x-axis (time) for better visualisation. **E)** Analysis of the data shown in (C). Particle tracking analysis was performed using the TrackMate plugin in Fiji to measure the total distance travelled and total displacement of PGC granules in *IP6K1^+/+^*and *IP6K1^-/-^* cells following carbachol stimulation over a 16-minute period. 850 particles were tracked in *IP6K1^+/+^* cells and 800 particles in *IP6K1^-/-^* cells, captured from four cells of each genotype. Data in (B) and (D) were analysed using a two-tailed Student’s t-test *(****P ≤ 0.0001; ***P≤ 0.0005;**P≤ 0.005)*.

### IP6K1 acts via SDC4 to support PGC granule dynamics

As active and catalytically inactive IP6K1 can restore PGC granule levels in IP6K1 depleted AGS cells, it is likely that IP6K1 acts via an interacting protein to support PGC granule dynamics. To identify the IP6K1 interactome in AGS cells we conducted tandem affinity purification of overexpressed SFB-tagged IP6K1, followed by mass spectrometry (Table S1). SFB-tagged GFP expressed in AGS cells was used as a control. We identified 90 proteins uniquely associated with IP6K1 that did not interact with the GFP control (Fig. 7A, Table S2). Among these were DDB1, UBE4A, and AP3B1, which are previously identified interactors of IP6K1 (49–51). The presence of these proteins in our IP6K1 pull-down validates the authenticity of our protein-protein interaction analysis. The localisation of IP6K1 to granules is unique to AGS cells, and was not observed in HEK293T cells (47). We reasoned that a protein which interacts with IP6K1 uniquely in AGS cells but not in HEK293T cells may underlie the function of IP6K1 in supporting granule dynamics. We therefore compared the list of IP6K1 interactors in these two cell lines, and identified 49 proteins that bind IP6K1 only in AGS cells (Fig. 7B, Table S3). Gene Ontology analysis to list biological processes associated with these proteins revealed that only two proteins, Annexin A2 (ANXA2), and Syndecan-4 (SDC4), are assigned GO terms related to the regulation of exocytosis (Table S4). We tested the localisation of GFP-tagged versions of these two proteins, and observed that ANXA2 is expressed throughout the cytoplasm, whereas SDC4 is localised to the plasma membrane and also to cytoplasmic granules in AGS cells (Fig. S4A). We therefore pursued SDC4 as a possible mediator of the effect of IP6K1 on granule dynamics. SDC4 is a member of the syndecan family of proteoglycans, which are composed of a variable N-terminal ectodomain, a single transmembrane domain, and a conserved cytoplasmic domain (42, 52). Syndecans are implicated primarily as mediators of endocytosis (52), but are also involved in exocytotic pathways, primarily in exosome biogenesis (53). In addition, SDC1 has been shown to act as a sorting receptor for soluble lipoprotein lipase (LIPL), directing its trafficking from the trans-Golgi network to secretory vesicles (54). In this context, we first assessed the expression of SDC4 in mouse stomach, and noted that it is present in gastric lysates from both *Ip6k1^+/+^* and *Ip6k1^-/-^* mice (Fig. S4B, C). Next, we confirmed the interaction between SDC4 and IP6K1 by expressing tagged versions of both proteins in AGS cells. GST-IP6K1 showed robust binding to SDC4 tagged C-terminally with mCherry (Fig. 7C). Additionally, endogenous SDC4 was found to interact with endogenous IP6K1 in AGS cells, supporting a potential functional association between these proteins (Fig. S4D). To determine whether binding with SDC4 is unique to IP6K1, or also observed in other IP6K paralogues, we co-expressed N-terminally myc-tagged IP6K1, IP6K2 or IP6K3 with SDC4 in AGS cells. Interestingly, the interaction between SDC4 and IP6K1 was distinct and not observed with IP6K2 or IP6K3 (Fig. 7D).

**Figure 7.**
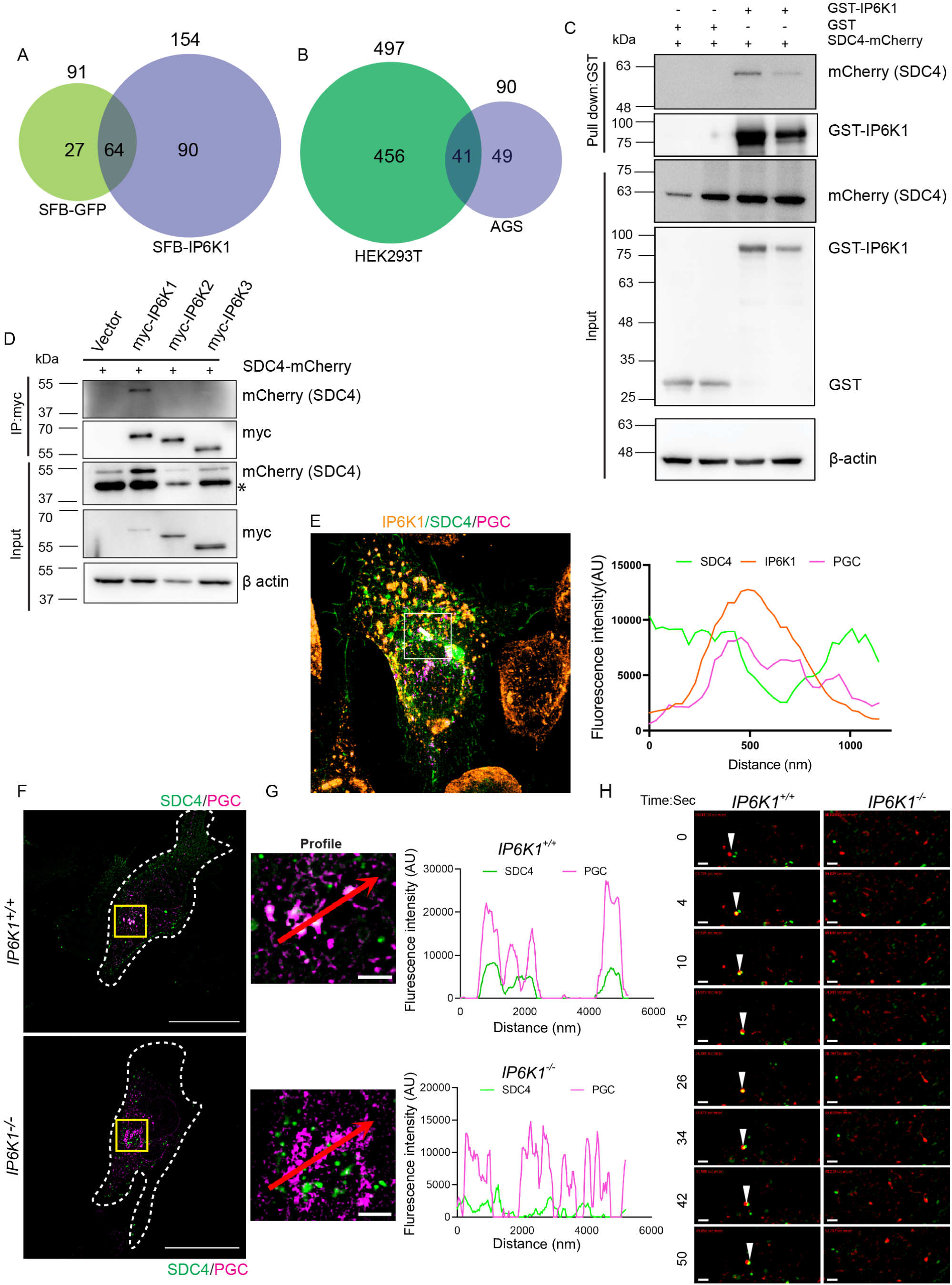
SDC4 mediates the effect of IP6K1 on PGC granule dynamics. **(A)** AGS cells were transfected to express either SFB-tagged IP6K1 or SFB-GFP as a control. SFB-tagged proteins were purified by tandem-affinity purification using streptavidin beads followed by S-protein beads in two replicates for each set. The Venn diagram illustrates the total number of interacting proteins identified across both replicates for each condition, with the numbers of GFP- and IP6K1-binding proteins indicated above. **B)** The Venn diagram illustrates IP6K1 interacting proteins identified earlier in HEK293T cells (51), in comparison with IP6K1 interactors in AGS cells identified in this study. **C)** GST-IP6K1 or GST alone were transiently overexpressed in wild-type AGS cells with SDC4-mCherry, and pulled down on glutathione agarose beads. The GST tag was detected using an anti-GST antibody to confirm the expression and successful pull-down of GST-tagged proteins. The experimental replicate data (1 and 2) shown here are representative of three independent experiments (N=3). **D)** Representative immunoblots show the co-immunoprecipitation of overexpressed SDC4-mCherry with myc-tagged IP6K paralogues. AGS cells were transiently transfected with plasmids encoding myc-IP6K1, myc-IP6K2, or myc-IP6K3. IP6Ks were immunoprecipitated and probed for SDC4-mCherry using an anti-mCherry rabbit polyclonal antibody. myc-tagged IP6Ks were detected using an anti-myc mouse monoclonal antibody (N=3). Non-specific bands in the input sample are marked with an asterisk (*). **E)** The image shows triple staining of fixed AGS cells for IP6K1-V5 (orange), SDC4-GFP (green), and PGC-mCherry (magenta). The corresponding intensity profile confirms partial co-localisation of these three proteins within the cell. **F)** Co-expression of SDC4-GFP (green) with PGC-mCherry (magenta) in *IP6K1^+/+^* and *IP6K1^-/-^* AGS cells. Dashed white lines mark the cell boundaries Scale bar 10 µm. **G)** Zoomed in images of the region in (F) marked by yellow boxes. Scale bar 2 μm. The traces display the fluorescence intensity profile for SDC4 (green) and PGC (magenta), measured along the red arrow in the figure. **H)** Live cell imaging was used to evaluate the co-migration of SDC4 and PGC in *IP6K1^+/+^* cells compared to *IP6K1^-/-^*AGS cells. Still frames captured within 60 seconds indicate co-migration of SDC4 (green) and PGC (red) granules, marked by white arrowheads in *IP6K1*^+/+^ cells. Scale bar: 1 μm. Super-resolution images in (E, F,G and H) were obtained with the Elyra 7 (SIM) module on a Zeiss LSM 980 confocal microscope, using a 63×/1.4 NA objective.

To probe whether SDC4 could be the link between IP6K1 and PGC granule assembly, we first examined whether SDC4 co-localises with PGC and IP6K1 in AGS cells. We observed partial co-localisation of these three proteins in some granules (Fig. 7E), confirming the association of IP6K1 with PGC and SDC4. Next, we examined whether IP6K1 affects SDC4 and PGC granule co-localisation. Whereas SDC4 and PGC showed partial co-localisation in *IP6K1^+/+^* cells, we did not see any co-localisation of these two proteins in *IP6K1^-/-^* AGS cells (Fig. 7F and G). Finally, we conducted live imaging of SDC4-GFP and PGC-mCherry expressed in *IP6K1^+/+^* and *IP6K1^-/-^* AGS cells (Video S3, S4). Still frames taken at different time intervals revealed co-migration of PGC and SDC4 in *IP6K1^+/+^* cells, whereas *IP6K1^-/-^* AGS cells failed to display either co-localization or co-migration of PGC and SDC4 (Fig. 7H). These observations suggest that IP6K1 and SDC4 act in conjunction to facilitate PGC granule assembly and secretion. Mis-localisation of SDC4 in the absence of IP6K1 likely results in reduced incorporation of PGC into secretory granules.

## DISCUSSION

IP6K1 is expressed in many mammalian tissues (1, 31–33), with highest expression levels recorded in the brain, testis, and pancreas, where it has been shown to participate in neuronal development and exocytosis, spermatogenesis, and insulin secretion respectively (10, 13, 17–19, 21, 24, 55). Additionally, IP6K1 expressed in muscle, liver and adipose tissue participates in metabolic homeostasis (22, 23, 25, 43, 56), and IP6K1 in the kidney supports renal tubular function (57, 58). Our study shows that IP6K1 is expressed throughout the mouse gastrointestinal tract, and is the first to address the function of IP6K1 in this organ system. In the stomach, IP6K1 expression was predominantly localised to granules in the cytoplasm of gastric chief cells, the secretory cells responsible for the production and release of the digestive enzymes PGC and LIPF. Loss of IP6K1 in mice disrupted the accumulation of these enzymes in secretory granules, impairing their release into the gastric lumen. The decrease in gastric pepsin in *Ip6k1^-/-^* mice correlated with increased faecal protein content, reduced muscle mass, and a sustained reduction in body weight, possibly indicative of impaired digestion in these mice. Using the AGS gastric adenocarcinoma cell line as a model, we found that IP6K1 is essential for secretory granule assembly and dynamics. IP6K1 localized to PGC-containing granules in AGS cells, a pattern absent in other cell lines including HEK293T and U-2 OS cells in which IP6K1 expression has been characterised (47). The depletion of IP6K1 in AGS cells disrupted PGC granule assembly, similar to the phenotype observed in gastric chief cells in *Ip6k1^-/-^* mice. This defect was rescued by both catalytically active and inactive IP6K1, indicating a 5-InsP_7_ synthesis-independent function for IP6K1 in PGC granule assembly. In addition, the loss of IP6K1 also impaired PGC granule dynamics and exocytosis upon stimulation of AGS cells with the acetylcholine mimic carbachol. A protein interactome analysis for IP6K1 in AGS cells revealed that it binds the transmembrane proteoglycan SDC4. In the absence of IP6K1, the co-localization and co-migration of SDC4 with PGC granules is disrupted in AGS cells, providing an explanation for the reduced formation of digestive enzyme granules observed in *Ip6k1*^-/-^ gastric chief cells (Fig. 8).

**Figure 8:**
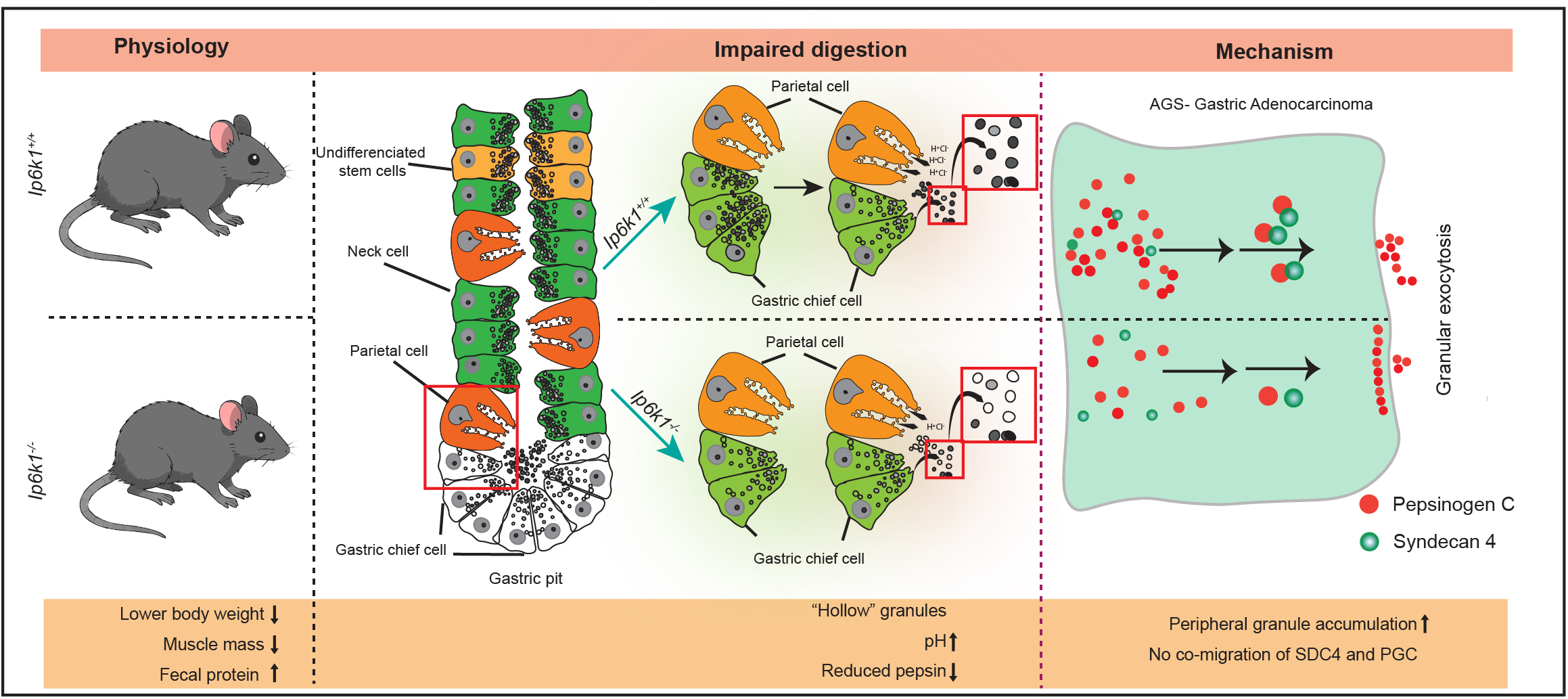
Schematic overview of IP6K1 function in gastric digestion. Illustration highlights the role of inositol phosphate kinase 1 (IP6K1) in regulating gastric digestion. Deletion of IP6K1 reduces serum protein and increases faecal protein, correlating with muscle loss and reduced body weight. Different cell types are marked in a representation of a gastric gland. The model highlights IP6K1’s role in gastric digestion through PGC and LIPF granule formation and secretion in chief cells. Finally, it shows that IP6K1 acts via SDC4 to facilitate PGC granule trafficking and release.

Our study revealed an exaggerated difference in body weight of juvenile *Ip6k1*^-/-^ mice compared with *Ip6k1*^+/+^ mice during the period of their rapid growth from ages 28 to 40 dpp (Fig. 1A). Earlier studies on the impact of IP6K1 on body weight have focused on adult mice or aged mice, with an emphasis on mechanisms by which the loss of IP6K1 protects mice from high fat diet-induced or age-dependent obesity (16, 22, 25, 43, 59). 5-InsP_7_ acts as an inhibitor of the Ser/Thr kinase AKT by competing with PtdIns(3,4,5)P_3_ for binding to the PH domain of AKT and preventing its translocation to the plasma membrane (22). Therefore, in *Ip6k1*^-/-^ mice, the reduction in 5-InsP_7_ results in hyperactivation of AKT in muscle, liver and white adipose tissue, which leads to increased insulin sensitivity and resistance to weight gain (22, 25, 60). In addition, the deletion of IP6K1 enhances thermogenic energy expenditure and increases lipolysis in adipocytes, also contributing to protection against high-fat diet induced obesity (43, 56, 59). Our study reveals a different mechanism by which the loss of IP6K1 may contribute to reduced body weight in mice. Impaired release of PGC and LIPF in the stomach of *Ip6k1*^-/-^ mice is likely to reduce protein and lipid digestion, contributing to reduced weight gain on a normal diet. Assessing the effect of impaired gastric lipase on triglyceride accumulation in adipose tissue would be confounded by other known actions of IP6K1 on lipid metabolism in adipocytes (22, 43, 56). We therefore only examined potential whole-body effects of impaired gastric protein digestion in *Ip6k1*^-/-^ mice by monitoring skeletal muscle mass, which we observed to be reduced in 2–3-month-old *Ip6k1*^-/-^ mice compared with their *Ip6k1*^+/+^ counterparts. This data is in contrast to an earlier observation that 10 month old *Ip6k1*^-/-^ mice show an increase in the mass of the gastrocnemius muscle compared with *Ip6k1*^+/+^ mice, correlating with an increase in the level of muscle glycogen accumulation (22). Changes in protein and carbohydrate metabolism as a function of age may underlie this observed divergence in the impact of IP6K1 deletion on skeletal muscle mass.

In exocrine cells, secretory granules bud from the trans-Golgi network, undergo maturation and homotypic fusion directed by Rabs, notably RAB26 and RAB3D in the case of zymogenic granules in gastric chief cells, to yield mature granules that are ready for exocytosis in response to secretagogues (61–65). Our discovery that the absence of IP6K1 impairs the biogenesis and release of PGC containing secretory granules, must be viewed in the context of other documented functions of IP6K1 in vesicle trafficking and exocytosis. 5-InsP_7_ synthesised by IP6K1 pyrophosphorylates the intermediate chain of the dynein motor protein complex, enhancing its interaction with dynactin to upregulate dynein-dependent transport of vesicles towards the minus-end of microtubules, so that the loss of IP6K1 in fibroblasts results in a fragmented Golgi architecture (8). Similar defects in Golgi architecture in mouse gastric chief cells and AGS cells deprived of IP6K1, which have been observed in the present study (Fig. S2D, E; S3A, B), may arise from the same molecular dysfunction, and contribute to impaired secretory granule biogenesis in these cells. Several studies have shown that IP6K1 regulates vesicle exocytosis in neurons and pancreatic β cells. IP6K1 acts independently of its catalytic activity to bind GRAB, a guanine nucleotide exchange factor (GEF) that activates RAB3A, which in turn inhibits synaptic vesicle exocytosis (66). IP6K1 binding with GRAB reduces RAB3A activity, leading to increased neurotransmitter release. Conversely, 5-InsP_7_ synthesised by IP6K1 has been shown to downregulate synaptic vesicle exocytosis in neurons by binding with synaptotagmin 1 (SYT1) to inhibit its fusogenic activity, so that knockdown of IP6K1 expression increased synaptic vesicle fusion with the neuronal plasma membrane (10, 11). The loss of IP6K1 has also been shown to increase exocytosis of the readily releasable pool of VGLUT1 and VGLUT2 containing synaptic vesicles in neurons (67). Although synaptic vesicles and secretory granules share many components, there are considerable differences in the mechanisms of their biogenesis and regulated exocytosis (61, 64, 68). In contrast to the predominantly down-regulatory impact of IP6K1 on synaptic vesicle exocytosis, IP6K1 strongly supports exocytosis of the readily releasable pool of insulin containing secretory granules in pancreatic β cells, acting via a mechanism involving the binding of synaptotagmin 7 (SYT7) with 5-InsP_7_ (20, 21, 24). Consequently, the loss of IP6K1 in mice leads to reduced serum insulin levels and impaired glucose stimulated insulin secretion in pancreatic β cells (16, 22, 24). A similar mechanism may underlie the decrease in PGC granule exocytosis that we observed in *IP6K1^-/-^*AGS cells (Fig. 6A, B).

In conclusion, our study identifies, for the first time, a functional role for IP6K1 in digestion physiology. We show that IP6K1 is needed for the proper assembly and release of digestive enzyme granules in gastric chief cells, a process central to efficient protein and lipid digestion in mammals. At the molecular level we have identified SDC4 as a novel IP6K1 interactor, which mediates IP6K1-driven PGC granule assembly. Our findings set the stage for future investigations into the role of IP6K1 in gastrointestinal tract physiology, and mechanisms of secretory granule biogenesis and release.

## MATERIALS AND METHODS

### Mice

All animal studies were performed in accordance with guidelines provided by the Committee for the Control and Supervision of Experiments on Animals, Ministry of Fisheries, Animal Husbandry and Dairying, Government of India, with protocols approved by the Institutional Animal Ethics Committee (Protocol Number EAF/RB/01/2025). The *Ip6k1* gene knockout mouse, carrying a deletion spanning the splice site, coding region and a portion of the 3′ UTR of the terminal exon (exon 6) of *Ip6k1*, was generated as described (16), and back-crossed into the C57BL/6J strain. Mice used in this study were housed in the Experimental Animal Facility at the Centre for DNA Fingerprinting and Diagnostics (CDFD), Hyderabad. *Ip6k1^+/+^*and *Ip6k1^−/−^* mice used for experimental analyses were generated by breeding *Ip6k1^+/-^* mice, and genotyping the offspring as described earlier (16). The experiments performed in this study required mice to be anaesthetised using isoflurane inhalation. Both male and female mice were used for the study. Gastrointestinal tract tissues were isolated from 2-3-month-old *Ip6k1^+/+^* and *Ip6k1^−/−^* mice after euthanizing them by CO_2_ inhalation. Body weight analysis and blood parameters were performed at different age spans as indicated in the figure legends.

### Reagents and antibodies

All chemicals were procured from Merck, unless specified otherwise. Primary antibodies used for immunoblotting (IB), immunofluorescence (IF), immunoprecipitation (IP), along with their dilutions, were as follows: anti-IP6K1 (Santa Cruz Biotechnology, sc-376290; 1:1000 IB; Sigma, HPA040825; 1:400 IF); anti-pepsinogen C (Abcam, ab180709; 1:5000 IB; 1:2000 IF); anti Lipase F (Sigma, HPA045930; 1:5000 IB;1:500 IF); anti E-cadherin (CST, 14472S; 1:500 IF); anti α-tubulin (Sigma-Aldrich, T9025; 1:5000 IB); anti β-actin (Sigma-Aldrich A2228; 1:5000 IB); anti SDC4 (Santa Cruz Biotechnology, sc-12766; 1:50 IF; 1:1000 IB); anti-GFP (Invitrogen, A11122; 1:5000 IB; 2 μg IP); anti-GST (Abcam, ab19256; 1:3000 IB), anti-cMyc (Sigma-Aldrich, M4439; 1:10,000 IB; 1 μg IP; 1:800 IF); anti mCherry (Abcam, ab183628; 2μg IP; 1:5000 IB).

### Serum protein and lipid profiling

Basal metabolic serum parameters including total serum protein, protein A/G ratio, lipid profile, blood urea nitrogen, and creatinine were examined in adult (2-3 month old) *Ip6k1^+/+^ and Ip6k1^-/-^*mice maintained on ad libitum food and water. Collected serum samples were stored at −20 °C and subsequently analysed at Rodenta Bioserve Labs, Hyderabad, using a Celltac α Hematology Analyzer (MEK-6550J/K; Nihon Kohden India Pvt. Ltd.) and a URA Semi-Automated Analyzer (Medsource Ozone Biomedicals Pvt. Ltd.).

### Measurement of gastrocnemius muscle mass in mice

In adult *Ip6k1^+/+^* and *Ip6k1^-/-^* mice, gastrocnemius muscle mass was evaluated relative to body weight. Animals were euthanized by CO₂ inhalation. To visualize the musculature, the skin was carefully removed, exposing the dorsal gastrocnemius and the ventral tibialis anterior muscles (69, 70). The gastrocnemius muscle from both lower limbs was then identified, dissected, and the wet muscle weight was recorded using a digital balance. The average gastrocnemius muscle mass for each mouse was divided by the total body weight of the mouse.

### Histology

Tissues of the gastrointestinal tract in *Ip6k1^+/+^* and *Ip6k1*^−/−^ mice, including stomach, duodenum, jejunum, ileum, colon and rectum, were subjected to histology. Mice were euthanized by CO2 inhalation, and dissected tissues were washed in phosphate buffer saline (PBS) and fixed in 10% formalin or in Bouin’s solution for 48 hours at 4°C. The tissues were then dehydrated in sequential ethanol (50%, 75%, 95% and 100%) and xylene to remove fixatives, and embedded in paraffin wax. 4 µm thick formalin-fixed paraffin-embedded (FFPE) sections were prepared on glass slides, deparaffinised by heating at 60°C for 1 h, cleared in xylene, and rehydrated in a graded series of ethanol (100%, 95%, 70%, and 50%). The sections were then stained using haematoxylin and eosin (H&E). Images were captured using a bright-field light microscope (Nikon ECLIPSE Ni-U, NIS Elements acquisition software; 20X 0.5NA, or 40X 0.75NA). H&E sections from age-matched *Ip6k1^+/+^* and *Ip6k1^-/-^* mouse tissues were subjected to detailed examination for any histopathological anomalies. Stomach tissue sections were analysed to determine the total number of gastric chief cells per gland in the fundus. The number of gastric chief cells per gastric gland was counted and compared in *Ip6k1^-/-^* and *Ip6k1^+/+^* stomach sections.

### Immunofluorescence of stomach tissue sections

4µm thick stomach sections were deparaffinized at 60°C for 1 h, cleared in xylene, and sequentially rehydrated with ethanol (100%, 95%, 70%, and 50%) and distilled water. Antigen retrieval was carried out in 10 mM sodium citrate buffer (pH 6.0) supplemented with 0.5% Tween-20. Following retrieval, tissues were permeabilized with 0.5% Triton X-100 for 15 min and subsequently blocked with either 4% FBS in PBST or 5% BSA in PBST. Primary antibody incubation was done at 4°C overnight and Alexa Fluor 488 (1:500) or Alexa Fluor 568 (1:500) goat anti-rabbit or anti-mouse IgG were incubated for 1 h in the dark at room temperature (RT). Slides were mounted using VectaShield with DAPI for nuclear staining. Images were captured on a Zeiss LSM 700 confocal microscope equipped with 405, 488 and 555 nm lasers, and fitted with a 63×1.4 NA objective.

### Lipase activity measurement

The dissected stomach was collected from *Ip6k1^+/+^* and *Ip6k1^-/-^* mice following euthanasia by CO_2_ inhalation. The fundus region of the stomach was isolated and rinsed in cold PBS. ∼40 mg tissue was placed in 100 µL assay solution supplied with the colorimetric Lipase Assay Kit (Abcam, ab102524). Homogenization was performed on ice using a Dounce homogenizer with 25 passes. The homogenates were centrifuged at 18,000 x g for 10 min at 4°C in a refrigerated centrifuge to remove insoluble material. The supernatant was transferred to a fresh tube and maintained on ice until further analysis. Samples were diluted 1:1 in assay buffer and were assayed in duplicates for lipase activity, following the manufacturer’s instructions, using different concentrations of glycerol to prepare the standard curve. Absorbance was recorded at 570 nm on a microplate reader (Perkin Elmer Enspire Multimode Plate Reader) in kinetic mode, every 3 minutes, for 60 min at 37°C protected from light. Lipase activity in the samples was extrapolated from the standard curve.

### Pyloric ligation for gastric secretion

Before conducting pyloric ligation, mice were fasted for 14-16 h, and transferred to clean cages with no bedding and ad libitum water. Mice were anaesthetised using an isoflurane inhalation chamber (1:1 oxygen:isoflurane) for 30 sec and immediately transferred to a thermal pad maintained at 37°C. For the duration of the pyloric ligation procedure, mice were kept anesthetized by administering 80% oxygen and 20% isoflurane as inhalation. The ventral side of the mice was sterilized with 70% ethanol and the abdominal hair was removed. 1 cm x 1 cm incision was made in the xiphoid cavity to access the peritoneum membrane, and a diagonal cut was made in the peritoneum membrane. To access the pyloric region, the stomach of the mice was gently mobilized out through the incision. After ligating the pyloric sphincter with sterile silk thread, the stomach was carefully repositioned into the abdominal cavity. The abdominal incision was sealed, and mice were allowed to recover until fully conscious and mobile. Following recovery from anaesthesia, subcutaneous injections of histamine (Sigma-Aldrich, 59964; 10 mg/kg body weight), carbachol (Sigma-Aldrich, Y0000113; 50 mg/kg body weight), or PBS were administered. At 1.5 h after stimulation, mice were euthanized, and the ligated stomach was carefully dissected to collect gastric secretions. Following pyloric ligation, the pH of the gastric fluid was measured using a handheld, small-volume digital pH meter (LAQUAact PH110, HORIBA) equipped with a Micro ToupH electrode (9618S-10D). Following pH measurement, gastric secretions were subjected to western blot analysis to assess pepsin levels.

### Plasmids

Mouse IP6K1 (GenBank ID: NM_013785.2) was cloned into the pMH-SFB vector to generate SFB-mIP6K1 using Gateway cloning (Invitrogen). SFB-GFP, pEGFP-N1 and pEGFP-C1 vectors were used as controls. The catalytically inactive mIP6K1 mutant (K226A) was generated by site-directed mutagenesis with overlap extension PCR and cloned in a similar manner. Untagged human Pepsinogen II (PGC) cDNA (HG12072-UT) in the pCMV3 vector was obtained from Sino Biological, and subcloned into the pcDNA3.1-mCherry backbone to generate PGC-mCherry. EYFP-Golgi was obtained from Clontech. Mouse SDC4 cDNA (GenBank ID: NM_011521.2) was subcloned into pEGFP-C1 and pcDNA3.1-mCherry to generate C-terminally tagged constructs, mSDC4-GFP and mSDC4-mCherry, respectively. The generation of C-terminally V5-tagged human IP6K1 (GenBank ID: NM_001242829.2) has been described earlier (15). Plasmids expressing myc-tagged mouse IP6K1 (GenBank ID: NM_013785.2), human IP6K2 (GenBank ID: NM_001005909.3), and human IP6K3 (GenBank ID: NM_001142883.2) in the pCMV-Myc-N backbone, as well as GST-mIP6K1 in pcDNA3.1-GST and pcDNA3.1-GST control plasmid, were kindly provided by Dr. Solomon Snyder (Johns Hopkins University School of Medicine, Baltimore, USA).

### Cell line and transfection

The human gastric carcinoma cell line AGS was obtained from the American type culture collection (ATCC; Manassas, VA, USA). AGS cells were cultured in RPMI 1640 supplemented with 10% FBS, 100 U/mL penicillin G sodium, 100 mg/mL streptomycin sulfate, and 1 mM L-glutamine, at 37°C and 5% CO2. Cells were transfected using polyethylenimine (1:3 DNA:PEI), and harvested 36 h post-transfection. The interactome of IP6K1 in AGS cells was determined by using tandem affinity purification of SFB-tagged IP6K1 from AGS cells, and SFB-GFP was used as a control. Briefly, AGS cells transiently expressing SFB-IP6K1 or SFB-GFP were lysed with NETN buffer (50 mM Tris-HCl pH 8.0, 150 mM NaCl, 1 mM EDTA and 0.5% Nonidet P-40) containing protease inhibitors and phosphatase inhibitors, for 1 h at 4°C. Lysates were then incubated with streptavidin-sepharose for 2 h at 4°C. After removing the unbound proteins by washing the beads three times with lysis buffer, the associated proteins were eluted using 2 mg/ml biotin (Merck) for 2 h at 4°C. The eluate was then incubated with S-protein agarose (Novagen) beads for 1 h at 4°C. After clearing the unbound proteins by washing, the proteins associated with S-protein agarose beads were eluted by boiling in 2x Laemmli buffer for 5 min at 95°C. The proteins were identified by mass spectrometry analysis carried out by the Taplin Biological Mass Spectrometry Facility (TMSF) at Harvard University.

### Generation of IP6K1 knockout AGS cell line

AGS knockout cells were generated using CRISPR/Cas9-mediated gene deletion. AGS cells were transfected with three sgRNAs targeting IP6K1 (sgRNA1: 5’ CCTTGGACTCGGAGCGCATG 3’; sgRNA2: 5’ GCTGCTGCTTGTGACAACGC 3’; sgRNA3: 5’ GAGCTTTCGGTCCTTGGACT 3’) cloned into the pU6-2A-GFP-2A-Puro plasmid. Following puromycin selection (1 µg/mL) for 5 days, cells were seeded by serial dilution in 96-well plates to isolate single colonies. Non-targeting sgRNA (5’ CTTACCCCTATTATAATGAA 3’) was used to generate a non-targeted control (IP6K1^+/+^) AGS cell line. Clones were genotyped to identify frameshift mutations in both alleles of IP6K1 as described earlier (47), and knockout (*IP6K1^-/-^*) was confirmed by western blotting and immunofluorescence.

### Immunofluorescence assay in AGS cells

AGS non-targeted control (*IP6K1^+/+^*) and *IP6K1^-/-^* cells were seeded on 12 mm glass coverslips in a 6-well plate, 12-well plate, or 35 mm dish. Cells were washed with PBS, fixed with 4% paraformaldehyde for 15 min, and permeabilised using PBS containing 0.15% Triton X-100 (PBST) for 10-12 min min at RT. Cells were incubated in blocking buffer (5% BSA or 4% FBS in PBST) for 1 h at RT, and overnight at 4°C with primary antibodies diluted in blocking solution. Following a triple rinsing with PBST, cells were subjected to 1 h incubation at RT in the dark with fluorophore-conjugated secondary antibodies. Coverslips were washed three times with PBST and mounted on glass slides using VectaShield with DAPI for nuclear staining. Images were captured on a Zeiss LSM 700/900 confocal microscope equipped with 405, 488 and 555 nm lasers, and fitted with a 63×1.4 NA objective, or a Leica TCS SP8 confocal microscope equipped with 405, 488, 514, 561, and 633 nm lasers using a 63x 1.4 NA oil immersion objective, or Elyra 7 structured illumination microscope (SIM) module of the Zeiss LSM 980 confocal microscope, equipped with 405, 488, 561, and 642 nm lasers, and 63x 1.4 NA oil immersion objective. The exposure settings and other imaging parameters were identical for images of different cell types/treatment conditions in a single experiment. All images shown are maximum intensity projections (MIP) of a Z-stack. Images were subjected to contrast or level adjustment for improved visualization using LASX (Leica application suit X; Ver-3.4.2.18368) or ZEN black/blue 3.6 software.

### Live cell imaging of AGS cells

AGS non-targeted control (*IP6K1^+/+^*) and *IP6K1^-/-^*cells were seeded in glass bottom 35 mm dishes 12 h before transfection. Cells were co-transfected with plasmids encoding PGC-mCherry and YFP-Golgi, or PGC-mCherry and SDC4-GFP, as indicated. Cells were imaged 48 h post-transfection, following treatment with 100 µM carbachol where indicated. Time-series images of cells expressing PGC-mCherry and SDC4-GFP were acquired using a Zeiss LSM 980 confocal microscope with Elyra 7 SIM for super-resolution, equipped with 405, 488, 561, and 642 nm lasers and a 63x 1.4 NA oil immersion objective. Particle tracking analysis (distance and displacement) was conducted using TrackMate plugin, and kymographs were plotted using the Kymograph Builder in Fiji software.

### Immunoprecipitation

Confluent AGS cells were lysed for 4 h at 4°C in a buffer containing 50 mM Tris-HCl (pH 8.0), 150 mM NaCl, 1 mM EDTA, 1% Triton X-100, and supplemented with protease (Merck, P8340) and phosphatase (Merck, P5762) inhibitor cocktails. Lysates were clarified by centrifugation at 16,000 × g for 15 min at 4°C and pre-cleared by incubation with protein A- or protein G-Sepharose beads (pre-equilibrated in lysis buffer) for 1 h at 4°C. For immunoprecipitation, specific antibodies were added to pre-cleared lysates and incubated overnight at 4°C. Immune complexes were captured with protein A- or G-Sepharose beads, washed extensively with lysis buffer, and eluted by boiling in 2× Laemmli buffer. For SDC4 interaction studies, cells were co-transfected with GST-tagged IP6K1 and mCherry-SDC4 constructs, and lysed as described above. Lysates were incubated with glutathione-Sepharose beads (Cytiva 17075601) pre-equilibrated in lysis buffer at 4°C for 2 h. After washing with lysis buffer, bound proteins were eluted by boiling in 2× Laemmli buffer and analysed by western blotting.

### Statistical analysis

Statistical analyses and graphing were performed using GraphPad Prism 8.4. Densitometry of immunoblots from three independent experiments was measured using Fiji or ImageJ. Protein band intensities were normalized to their respective loading controls and expressed relative to the control in each blot, as detailed in the figure legends. Comparisons between *Ip6k1^+/+^* and *Ip6k1^-/-^* genotypes were made using a two-tailed unpaired Student’s t-test, with P ≤ 0.05 considered significant. Data presentation and statistical details are provided in the corresponding figure legends.

## AUTHOR CONTRIBUTIONS

J.S. and R.B conceived and designed research; J.S. performed experiments with some assistance from P.P.; J.S and R.B analyzed data; J.S. and R.B interpreted results of experiments; J.S. prepared figures; J.S. and R.B drafted manuscript; J.S. and R.B edited and revised manuscript; J.S., P.P., and R.B. approved final version of manuscript.

## ACKNOWLEDGEMENTS

We gratefully acknowledge Rupinder Kaur, Maddika Subba Reddy and P. Chandra Shekar for generously sharing reagents. We thank the following colleagues for generating plasmids used in this study - Sitalakshmi Thampatty for SFB-GFP and SFB-IP6K1; Vineesha Oddi for IP6K1-V5; and Shubhra Ganguli for SFB-IP6K1 K226A. We appreciate the support of the technical staff at the Sophisticated Equipment Facility (SEF) and Experimental Animal Facility (EAF) at CDFD. We also thank Maddika Subba Reddy, P. Chandra Shekar, and members of Lab of Cell Signalling for their valuable feedback.

## DATA AVAILABILITY STATEMENT

The mass spectrometry data from this study are available on **MassIVE**, a ProteomeXchange Consortium member, and can be accessed through the dataset identifier **MSV000092121** at https://massive.ucsd.edu.

---

**Figure S1:**
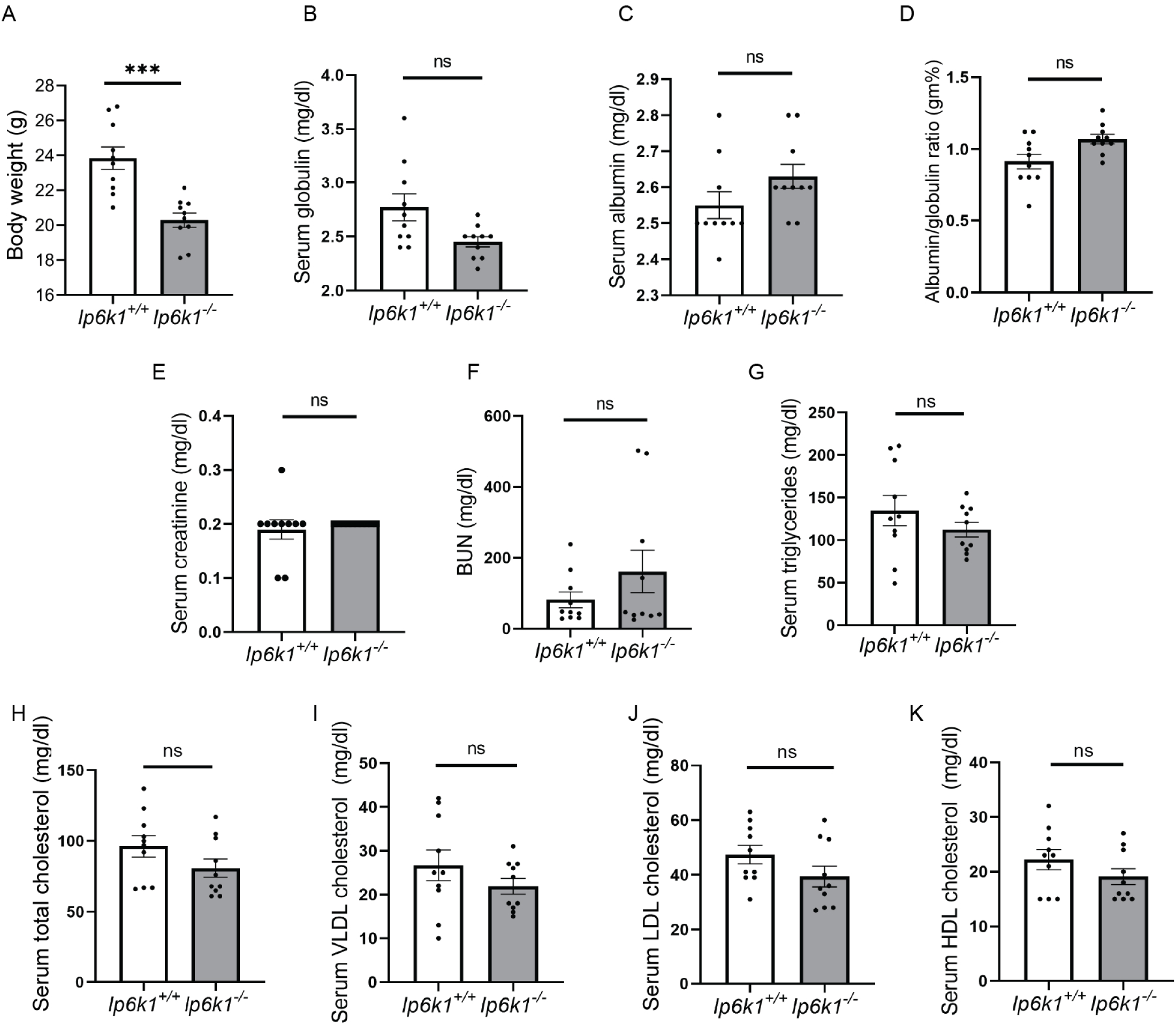
Impact of IP6K1 deletion on basal metabolic parameters in mice. **A-K)** Basal serum parameters at ad libitum food and water, to analyse serum protein and lipid profiles in adult *Ip6k1^-/-^* and *Ip6k1^+/+^*mice. The representative data for independent serum parameters in mice were analysed using a two-tailed unpaired Student’s t-test (mean ± SEM N=10 mice of each genotype; male and female combined) *(****P≤ 0.0005;* ns *P*≥ 0.05).

**Figure S2:**
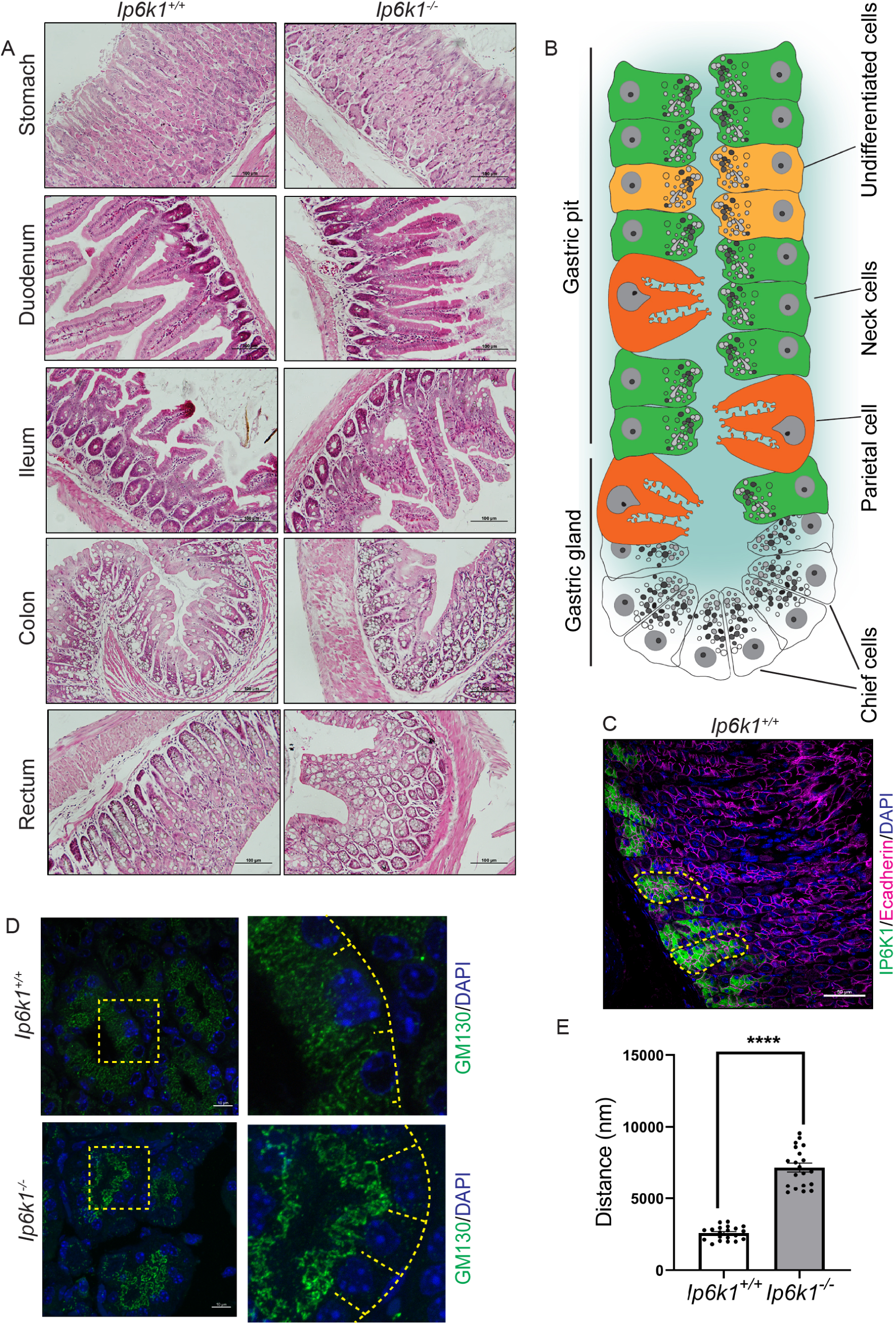
Histological analysis of gastrointestinal tract tissues of *Ip6k1^+/+^* and *Ip6k1^-/-^* mice. **A)** Haematoxylin and eosin-stained gastrointestinal tract tissue cross-sections of adult *Ip6k1^+/+^*and *Ip6k1^−/−^* mice. Histological analysis of stomach, duodenum, ileum, colon, and rectum, was performed to look for pathological changes or abnormal morphology. Tissues were examined for any changes across the mucosa, submucosa, muscularis propria, and serosa, as well as for any histological abnormalities such as inflammation, hyperplasia, hypertrophy, metaplasia, dysplasia, neoplasia, or hypertrophy. Scale bars, 100 μm**. B)** Schematic diagram of a gastric gland illustrating the spatial arrangement and types of cells within the gastric pits and gland. **C)** Representative 40X image of *Ip6k1^+/+^* FFPE stomach sections showing IP6K1 expression in gastric glands (marker by yellow dashed line), but not in other cell types. Sections were co-stained with E-cadherin (cell boundaries) and DAPI (nuclei). **D)** Immunofluorescence staining of GM130 (green) in stomach FFPE sections of *Ip6k1^+/+^* and *Ip6k1^-/-^* mice. Yellow boxed region in images on the left are zoomed in on the right. Dashed lines on images on the right mark the basal membranes of gastric chief cells (N=2 mice of each genotype). Images were captured on a Zeiss LSM700 confocal microscope using 63X/1.4 NA objective. **E)** Quantification of images in (D) measuring the distance from the basal membrane to the Golgi marked by GM130 in gastric chief cells (N=20 glands). Statistical significance was assessed using a Student’s t-test (*\*\*\*\*P≤0.0005*), revealing altered GM130 distribution indicative of abnormal Golgi structure in *Ip6k1*^-/-^ gastric chief cells.

**Figure S3:**
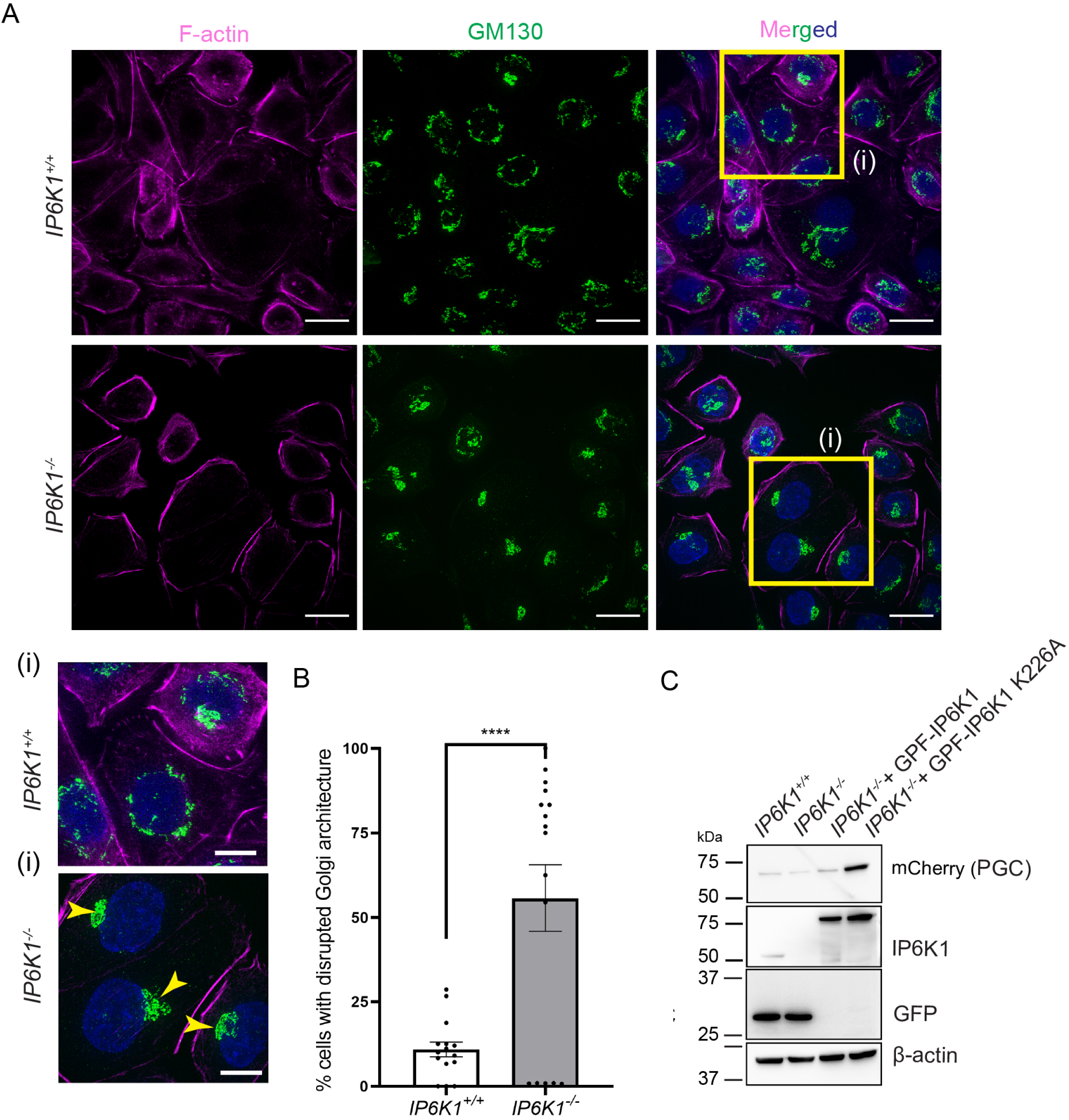
IP6K1 deletion disrupts Golgi architecture in AGS cells. **A)** Immunofluorescence analysis of AGS cells to detect the *cis*-Golgi marker-GM130 (green). F-actin was stained using rhodamine-phalloidin (magenta) to assist in identifying individual cells. Nuclei were stained with DAPI (blue). Disrupted Golgi morphology in *IP6K1^-/-^* AGS cells is indicated by yellow arrowheads in the inset (i). Scale bar 20 µm in (A) and 10 µm in (i). The images represent the maximum intensity projection (MIP) of z-stacks with xy dimensions displayed. Images were captured using an Elyra 7 (SIM) module of the Zeiss LSM 980 confocal microscope with 63×/1.4 NA Objective. **B)** Quantification of data in (A). The percentage of cells with disrupted Golgi morphology was calculated for each frame, which captured 17-22 cells. The graphs show mean±SEM of data from 14 frames for *IP6K1^+/+^* and 16 frames for *IP6K1^−/−^*cells, obtained over two independent experiments. The number of cells examined (n) were N=226 for IP6K1^+/+^ and N=228 for IP6K1^−/−^ cells. Data were analysed using a two-tailed unpaired Student’s t-test (\*\*\*\**P≤* 0.0001). **C)** Immunoblot showing loss of endogenous IP6K1 in *IP6K1^−/−^* AGS cells. Overexpression of GFP-IP6K1 (active), GFP-IP6K1 K226A (inactive), and PGC-mCherry was detected using anti-IP6K1 and anti-mCherry antibodies. β-actin was used as a loading control (N=2).

**Figure S4:**
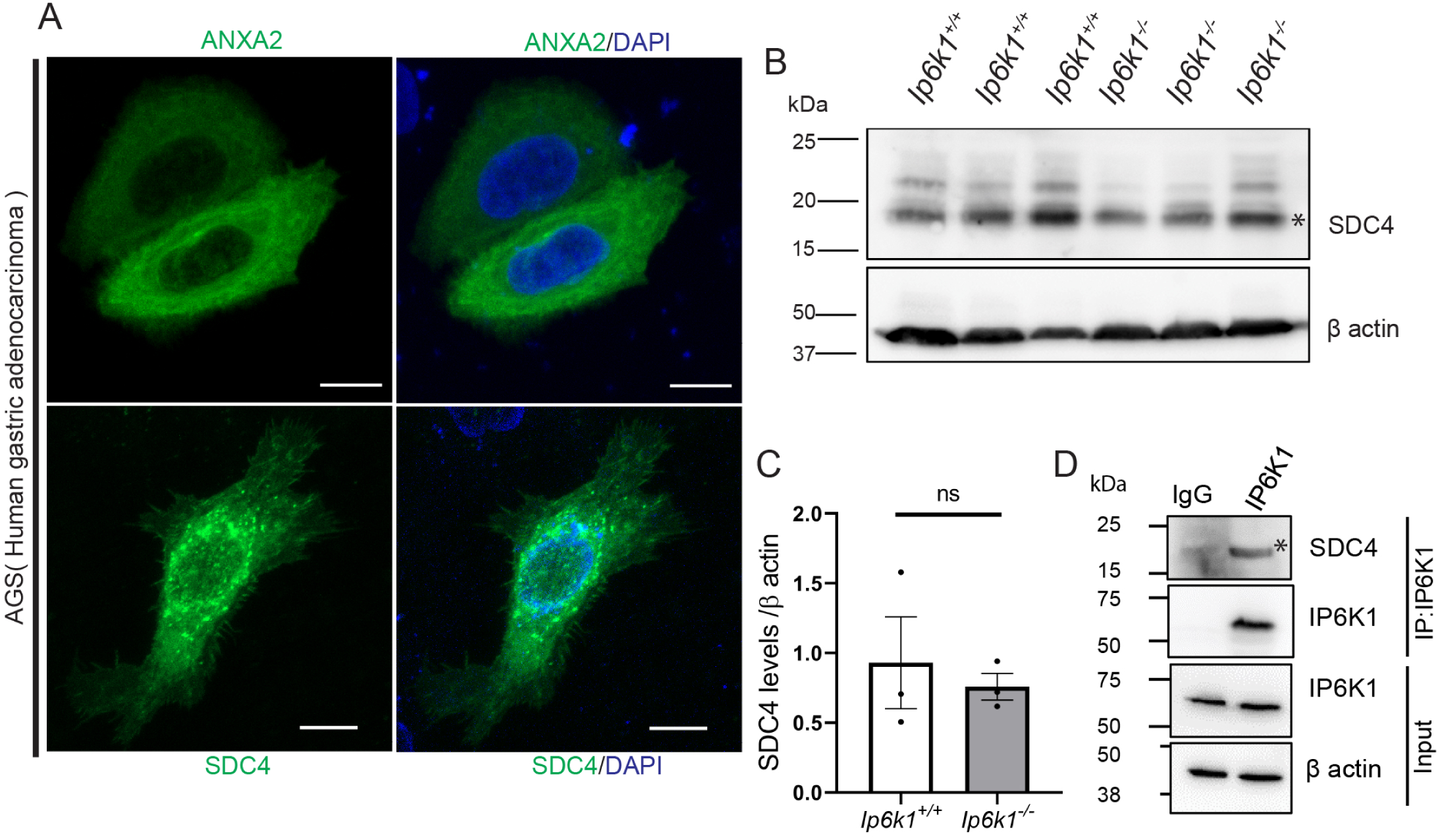
Endogenous interaction between SDC4 and IP6K1 in AGS cells. **A)** Immunofluorescence images of AGS cells show ANXA2 (green, top row) and SDC4 (green, bottom row) localization, with nuclei counterstained using DAPI (blue); both proteins display cytoplasmic and membrane-associated distribution (Scale bars 10 μm). **B)** Immunoblot analysis of SDC4 in stomach tissues of *Ip6k1^+/+^* mice and *Ip6k1*^−/−^ mice. β-actin was used as a loading control. The asterisk (*) indicates the specific band corresponding to SDC4 in the stomach tissues. SDC4 expression in stomach tissue was examined using the same lysates as shown in Fig. 3E for comparison. **C)** Quantification of SDC4 expression in adult *Ip6k1^+/+^* and *Ip6k1*^−/−^ mouse stomach. Values indicate mean ± SEM; N=3 for each genotype. Data were analysed using a two-tailed Student’s t-test (ns non-significant; *P*≥ 0.05). **D)** AGS cell extracts were immunoprecipitated using an antibody against the N-terminus of IP6K1, and normal rabbit IgG was used as a control. The presence of SDC4 was detected using an anti-SDC4 monoclonal antibody. β-actin was used as a loading control for input samples (N=3).

**Video S1, S2: PGC granule dynamics following carbachol stimulation in AGS cells.** Live imaging of PGC-mCherry and YFP-Golgi expressing *IP6K1*^+/+^ (S1) and *IP6K1*^-/-^ (S2) AGS cells following treatment with carbachol (100 µM). Images were captured at 1 image per 3.8 sec over a period of ∼16 min to obtain a total of 200 frames, using the Elyra 7 (SIM) module of the Zeiss LSM 980 confocal microscope with 63X/1.4 NA Objective. Images were converted to a video format at 5 frames per sec using Zen black software.

**Video S3, S4: PGC and SDC4 co-localisation and co-migration in AGS cells.** Live imaging of PGC-mCherry and SDC4-GFP expressing *IP6K1*^+/+^ (S3) and *IP6K1*^-/-^ (S4) AGS cells. Images were captured at 1 image per 3.8 sec to obtain a total of 100 frames, using the Elyra 7 (SIM) module of the Zeiss LSM 980 confocal microscope with 63X/1.4 NA Objective. Images were converted to a video format at 5 frames per sec using Zen black software.

**Table S1:** Mass spectrometry data showing counts and intensities for each protein in duplicate pull-downs of SFB-GFP and SFB-IP6K1 expressed in AGS cells.

**Table S2:** Comparison of SFB-GFP and SFB-IP6K1 interacting proteins identified in AGS cells.

**Table S3:** Comparison of SFB-IP6K1 binding proteins identified in HEK293T cells (51) and AGS cells.

**Table S4:** Analysis of 49 IP6K1 interacting proteins identified in AGS cells, that are absent in the IP6K1 interactome in HEK93T cells, using Enrichr tool to identify Biological Process GO terms (maayanlab.cloud/Enrichr).

## REFERENCES

1. Saiardi A, Erdjument-Bromage H, Snowman AM, Tempst P, and Snyder SH. Synthesis of diphosphoinositol pentakisphosphate by a newly identified family of higher inositol polyphosphate kinases. Current Biology 9: 1323–1326, 1999.

2. Thomas MP, and Potter BVL. The enzymes of human diphosphoinositol polyphosphate metabolism. The FEBS Journal 281: 14–33, 2014.

3. Shah A, Ganguli S, Sen J, and Bhandari R. Inositol Pyrophosphates: Energetic, Omnipresent and Versatile Signalling Molecules. J Indian Inst Sci 97: 23–40, 2017.

4. Saiardi A, Nagata E, Luo HR, Snowman AM, and Snyder SH. Identification and characterization of a novel inositol hexakisphosphate kinase. Journal of Biological Chemistry 276: 39179–39185, 2001.

5. Chakkour M, and Greenberg ML. Insights into the roles of inositol hexakisphosphate kinase 1 (IP6K1) in mammalian cellular processes. Journal of Biological Chemistry 300: 107116, 2024.

6. Jadav RS, Chanduri MV, Sengupta S, and Bhandari R. Inositol pyrophosphate synthesis by inositol hexakisphosphate kinase 1 is required for homologous recombination repair. J Biol Chem 288: 3312–3321, 2013.

7. Rao F, Xu J, Fu C, Cha JY, Gadalla MM, Xu R, Barrow JC, and Snyder SH. Inositol pyrophosphates promote tumor growth and metastasis by antagonizing liver kinase B1. Proc Natl Acad Sci U S A 112: 1773–1778, 2015.

8. Chanduri M, Rai A, Malla AB, Wu M, Fiedler D, Mallik R, and Bhandari R. Inositol hexakisphosphate kinase 1 (IP6K1) activity is required for cytoplasmic dynein-driven transport. Biochem J 473: 3031–3047, 2016.

9. Azevedo C, Burton A, Ruiz-Mateos E, Marsh M, and Saiardi A. Inositol pyrophosphate mediated pyrophosphorylation of AP3B1 regulates HIV-1 Gag release. Proc Natl Acad Sci U S A 106: 21161–21166, 2009.

10. Lee T-S, Lee J-Y, Kyung JW, Yang Y, Park SJ, Lee S, Pavlovic I, Kong B, Jho YS, Jessen HJ, Kweon D-H, Shin Y-K, Kim SH, Yoon T-Y, and Kim S. Inositol pyrophosphates inhibit synaptotagmin-dependent exocytosis. Proceedings of the National Academy of Sciences 113: 8314–8319, 2016.

11. Park SJ, Park H, Kim M-G, Zhang S, Park SE, Kim S, and Chung C. Inositol Pyrophosphate Metabolism Regulates Presynaptic Vesicle Cycling at Central Synapses. iScience 23: 101000, 2020.

12. Jadav RS, Kumar D, Buwa N, Ganguli S, Thampatty SR, Balasubramanian N, and Bhandari R. Deletion of inositol hexakisphosphate kinase 1 (IP6K1) reduces cell migration and invasion, conferring protection from aerodigestive tract carcinoma in mice. Cell Signal 28: 1124–1136, 2016.

13. Fu C, Xu J, Cheng W, Rojas T, Chin AC, Snowman AM, Harraz MM, and Snyder SH. Neuronal migration is mediated by inositol hexakisphosphate kinase 1 via α-actinin and focal adhesion kinase. Proceedings of the National Academy of Sciences 114: 2036–2041, 2017.

14. Burton A, Azevedo C, Andreassi C, Riccio A, and Saiardi A. Inositol pyrophosphates regulate JMJD2C-dependent histone demethylation. Proceedings of the National Academy of Sciences 110: 18970–18975, 2013.

15. Shah A, and Bhandari R. IP6K1 upregulates the formation of processing bodies by influencing protein-protein interactions on the mRNA cap. J Cell Sci 134: 2021.

16. Bhandari R, Juluri KR, Resnick AC, and Snyder SH. Gene deletion of inositol hexakisphosphate kinase 1 reveals inositol pyrophosphate regulation of insulin secretion, growth, and spermiogenesis. Proc Natl Acad Sci U S A 105: 2349–2353, 2008.

17. Malla AB, and Bhandari R. IP6K1 is essential for chromatoid body formation and temporal regulation of Tnp2 and Prm2 expression in mouse spermatids. J Cell Sci 130: 2854–2866, 2017.

18. Bhat SA, Malla AB, Oddi V, Sen J, and Bhandari R. Inositol hexakisphosphate kinase 1 is essential for cell junction integrity in the mouse seminiferous epithelium. Biochimica et Biophysica Acta (BBA) - Molecular Cell Research 1871: 119596, 2024.

19. Fu C, Rojas T, Chin AC, Cheng W, Bernstein IA, Albacarys LK, Wright WW, and Snyder SH. Multiple aspects of male germ cell development and interactions with Sertoli cells require inositol hexakisphosphate kinase-1. Scientific reports 8: 7039, 2018.

20. Illies C, Gromada J, Fiume R, Leibiger B, Yu J, Juhl K, Yang S-N, Barma DK, Falck JR, Saiardi A, Barker CJ, and Berggren P-O. Requirement of Inositol Pyrophosphates for Full Exocytotic Capacity in Pancreatic β Cells. Science 318: 1299–1302, 2007.

21. Rajasekaran SS, Kim J, Gaboardi GC, Gromada J, Shears SB, Dos Santos KT, Nolasco EL, Ferreira SS, Illies C, Köhler M, Gu C, Ryu SH, Martins JO, Darè E, Barker CJ, and Berggren PO. Inositol hexakisphosphate kinase 1 is a metabolic sensor in pancreatic β-cells. Cell Signal 46: 120–128, 2018.

22. Chakraborty A, Koldobskiy MA, Bello NT, Maxwell M, Potter JJ, Juluri KR, Maag D, Kim S, Huang AS, Dailey MJ, Saleh M, Snowman AM, Moran TH, Mezey E, and Snyder SH. Inositol pyrophosphates inhibit Akt signaling, thereby regulating insulin sensitivity and weight gain. Cell 143: 897–910, 2010.

23. Mukherjee S, Chakraborty M, Ulmasov B, McCommis K, Zhang J, Carpenter D, Msengi EN, Haubner J, Guo C, Pike DP, Ghoshal S, Ford DA, Neuschwander-Tetri BA, and Chakraborty A. Pleiotropic actions of IP6K1 mediate hepatic metabolic dysfunction to promote nonalcoholic fatty liver disease and steatohepatitis. Molecular Metabolism 54: 101364, 2021.

24. Zhang X, Li N, Zhang J, Zhang Y, Yang X, Luo Y, Zhang B, Xu Z, Zhu Z, Yang X, Yan Y, Lin B, Wang S, Chen D, Ye C, Ding Y, Lou M, Wu Q, Hou Z, Zhang K, Liang Z, Wei A, Wang B, Wang C, Jiang N, Zhang W, Xiao G, Ma C, Ren Y, Qi X, Han W, Wang C, and Rao F. 5-IP7 is a GPCR messenger mediating neural control of synaptotagmin-dependent insulin exocytosis and glucose homeostasis. Nature Metabolism 3: 1400–1414, 2021.

25. Ghoshal S, Mukherjee S, Chakraborty M, Msengi EN, Haubner J, and Chakraborty A. Whole Body Ip6k1 Deletion Protects Mice from Age-Induced Weight Gain, Insulin Resistance and Metabolic Dysfunction. Int J Mol Sci 23: 2022.

26. Ghosh S, Shukla D, Suman K, Lakshmi BJ, Manorama R, Kumar S, and Bhandari R. Inositol hexakisphosphate kinase 1 maintains hemostasis in mice by regulating platelet polyphosphate levels. Blood 122: 1478–1486, 2013.

27. Hou Q, Liu F, Chakraborty A, Jia Y, Prasad A, Yu H, Zhao L, Ye K, Snyder SH, and Xu Y. Inhibition of IP6K1 suppresses neutrophil-mediated pulmonary damage in bacterial pneumonia. Science translational medicine 10: eaal4045, 2018.

28. Lee H, Park SJ, Hong S, Lim SW, and Kim S. Deletion of IP6K1 in mice accelerates tumor growth by dysregulating the tumor-immune microenvironment. Anim Cells Syst (Seoul) 26: 19–27, 2022.

29. Kim MG, Zhang S, Park H, Park SJ, Kim S, and Chung C. Inositol hexakisphosphate kinase-1 is a key mediator of prepulse inhibition and short-term fear memory. Mol Brain 13: 72, 2020.

30. Heitmann T, and Barrow JC. The Role of Inositol Hexakisphosphate Kinase in the Central Nervous System. Biomolecules 13: 2023.

31. Pontén F, Jirström K, and Uhlen M. The Human Protein Atlas—a tool for pathology. The Journal of Pathology 216: 387–393, 2008.

32. Uhlén M, Fagerberg L, Hallström BM, Lindskog C, Oksvold P, Mardinoglu A, Sivertsson Å, Kampf C, Sjöstedt E, Asplund A, Olsson I, Edlund K, Lundberg E, Navani S, Szigyarto CA, Odeberg J, Djureinovic D, Takanen JO, Hober S, Alm T, Edqvist PH, Berling H, Tegel H, Mulder J, Rockberg J, Nilsson P, Schwenk JM, Hamsten M, von Feilitzen K, Forsberg M, Persson L, Johansson F, Zwahlen M, von Heijne G, Nielsen J, and Pontén F. Proteomics. Tissue-based map of the human proteome. Science 347: 1260419, 2015.

33. Papatheodorou I, Fonseca NA, Keays M, Tang YA, Barrera E, Bazant W, Burke M, Füllgrabe A, Fuentes AM, George N, Huerta L, Koskinen S, Mohammed S, Geniza M, Preece J, Jaiswal P, Jarnuczak AF, Huber W, Stegle O, Vizcaino JA, Brazma A, and Petryszak R. Expression Atlas: gene and protein expression across multiple studies and organisms. Nucleic Acids Res 46: D246–d251, 2018.

34. Betts JG, Kelly A. Young, James A. Wise, Eddie Johnson, Brandon Poe, Dean H. Kruse, Oksana Korol, Jody E. Johnson, Mark Womble, Peter DeSaix. Chapter 23.4 The Stomach. In: *Anatomy and Physiology*OpenStax, 2013, p. 1317.

35. Engevik AC, Kaji I, and Goldenring JR. The Physiology of the Gastric Parietal Cell. Physiological Reviews 100: 573–602, 2020.

36. Owen DA. Normal histology of the stomach. Am J Surg Pathol 10: 48–61, 1986.

37. Schubert ML. Gastric acid secretion. Current Opinion in Gastroenterology 32: 452–460, 2016.

38. Armand M. Lipases and lipolysis in the human digestive tract: where do we stand? Current Opinion in Clinical Nutrition & Metabolic Care 10: 156–164, 2007.

39. Lim SY, Steiner JM, and Cridge H. Lipases: it’s not just pancreatic lipase! Am J Vet Res 83: 2022.

40. Raufman JP. Gastric chief cells: receptors and signal-transduction mechanisms. Gastroenterology 102: 699–710, 1992.

41. Basque JR, and Ménard D. Establishment of culture systems of human gastric epithelium for the study of pepsinogen and gastric lipase synthesis and secretion. Microsc Res Tech 48: 293–302, 2000.

42. Elfenbein A, and Simons M. Syndecan-4 signaling at a glance. J Cell Sci 126: 3799–3804, 2013.

43. Zhu Q, Ghoshal S, Rodrigues A, Gao S, Asterian A, Kamenecka TM, Barrow JC, and Chakraborty A. Adipocyte-specific deletion of Ip6k1 reduces diet-induced obesity by enhancing AMPK-mediated thermogenesis. J Clin Invest 126: 4273–4288, 2016.

44. Giakoumaki I, Pollock N, Aljuaid T, Sannicandro AJ, Alameddine M, Owen E, Myrtziou I, Ozanne SE, Kanakis I, Goljanek-Whysall K, and Vasilaki A. Postnatal Protein Intake as a Determinant of Skeletal Muscle Structure and Function in Mice—A Pilot Study. International Journal of Molecular Sciences 23: 8815, 2022.

45. Aoyama S, Kim HK, Hirooka R, Tanaka M, Shimoda T, Chijiki H, Kojima S, Sasaki K, Takahashi K, Makino S, Takizawa M, Takahashi M, Tahara Y, Shimba S, Shinohara K, and Shibata S. Distribution of dietary protein intake in daily meals influences skeletal muscle hypertrophy via the muscle clock. Cell Rep 36: 109336, 2021.

46. Moreno P, Fexova S, George N, Manning JR, Miao Z, Mohammed S, Muñoz-Pomer A, Fullgrabe A, Bi Y, Bush N, Iqbal H, Kumbham U, Solovyev A, Zhao L, Prakash A, García-Seisdedos D, Kundu Deepti J, Wang S, Walzer M, Clarke L, Osumi-Sutherland D, Tello-Ruiz Marcela K, Kumari S, Ware D, Eliasova J, Arends Mark J, Nawijn Martijn C, Meyer K, Burdett T, Marioni J, Teichmann S, Vizcaíno Juan A, Brazma A, and Papatheodorou I. Expression Atlas update: gene and protein expression in multiple species. Nucleic Acids Research 50: D129–D140, 2021.

47. Morgan JAM, Singh A, Kurz L, Nadler-Holly M, Ruwolt M, Ganguli S, Sharma S, Penkert M, Krause E, Liu F, Bhandari R, and Fiedler D. Extensive protein pyrophosphorylation revealed in human cell lines. Nature Chemical Biology 2024.

48. Kim S, Bhandari R, Brearley CA, and Saiardi A. The inositol phosphate signalling network in physiology and disease. Trends Biochem Sci 49: 969–985, 2024.

49. Huttlin EL, Bruckner RJ, Navarrete-Perea J, Cannon JR, Baltier K, Gebreab F, Gygi MP, Thornock A, Zarraga G, Tam S, Szpyt J, Gassaway BM, Panov A, Parzen H, Fu S, Golbazi A, Maenpaa E, Stricker K, Guha Thakurta S, Zhang T, Rad R, Pan J, Nusinow DP, Paulo JA, Schweppe DK, Vaites LP, Harper JW, and Gygi SP. Dual proteome-scale networks reveal cell-specific remodeling of the human interactome. Cell 184: 3022–3040.e3028, 2021.

50. Rao F, Xu J, Khan AB, Gadalla MM, Cha JY, Xu R, Tyagi R, Dang Y, Chakraborty A, and Snyder SH. Inositol hexakisphosphate kinase-1 mediates assembly/disassembly of the CRL4-signalosome complex to regulate DNA repair and cell death. Proc Natl Acad Sci U S A 111: 16005–16010, 2014.

51. Hamid A, Ladke Jayashree S, Shah A, Ganguli S, Pal M, Singh A, and Bhandari R. Interaction with IP6K1 supports pyrophosphorylation of substrate proteins by the inositol pyrophosphate 5-InsP7. Bioscience Reports 44: 2024.

52. Afratis NA, Nikitovic D, Multhaupt HAB, Theocharis AD, Couchman JR, and Karamanos NK. Syndecans – key regulators of cell signaling and biological functions. The FEBS Journal 284: 27–41, 2017.

53. Baietti MF, Zhang Z, Mortier E, Melchior A, Degeest G, Geeraerts A, Ivarsson Y, Depoortere F, Coomans C, Vermeiren E, Zimmermann P, and David G. Syndecan– syntenin–ALIX regulates the biogenesis of exosomes. Nature Cell Biology 14: 677–685, 2012.

54. Sundberg EL, Deng Y, and Burd CG. Syndecan-1 Mediates Sorting of Soluble Lipoprotein Lipase with Sphingomyelin-Rich Membrane in the Golgi Apparatus. Dev Cell 51: 387–398.e384, 2019.

55. Illies C, Gromada J, Fiume R, Leibiger B, Yu J, Juhl K, Yang SN, Barma DK, Falck JR, Saiardi A, Barker CJ, and Berggren PO. Requirement of inositol pyrophosphates for full exocytotic capacity in pancreatic beta cells. Science 318: 1299–1302, 2007.

56. Ghoshal S, Tyagi R, Zhu Q, and Chakraborty A. Inositol hexakisphosphate kinase-1 interacts with perilipin1 to modulate lipolysis. The international journal of biochemistry & cell biology 78: 149–155, 2016.

57. Haykir B, Saiardi A, Hernando N, and Wagner C. Inositol hexakisphosphate kinases 1 and 2 (IP6K1/2) are involved in renal control of phosphate homeostasis. The FASEB Journal 35: 2021.

58. Gu C, Liu J, Liu X, Zhang H, Luo J, Wang H, Locasale JW, and Shears SB. Metabolic supervision by PPIP5K, an inositol pyrophosphate kinase/phosphatase, controls proliferation of the HCT116 tumor cell line. Proc Natl Acad Sci U S A 118: 2021.

59. Zhu Q, Ghoshal S, Tyagi R, and Chakraborty A. Global IP6K1 deletion enhances temperature modulated energy expenditure which reduces carbohydrate and fat induced weight gain. Molecular metabolism 6: 73–85, 2017.

60. Luo HR, Huang YE, Chen JC, Saiardi A, Iijima M, Ye K, Huang Y, Nagata E, Devreotes P, and Snyder SH. Inositol Pyrophosphates Mediate Chemotaxis in *Dictyostelium* via Pleckstrin Homology Domain-PtdIns(3,4,5)P3 Interactions. Cell 114: 559–572, 2003.

61. Burgoyne RD, and Morgan A. Secretory Granule Exocytosis. Physiological Reviews 83: 581–632, 2003.

62. Tian X, Jin RU, Bredemeyer AJ, Oates EJ, Błażewska KM, McKenna CE, and Mills JC. RAB26 and RAB3D Are Direct Transcriptional Targets of MIST1 That Regulate Exocrine Granule Maturation. Molecular and Cellular Biology 30: 1269–1284, 2010.

63. Kögel T, and Gerdes HH. Maturation of secretory granules. Results and problems in cell differentiation 50: 1–20, 2010.

64. Bonnemaison ML, Eipper BA, and Mains RE. Role of Adaptor Proteins in Secretory Granule Biogenesis and Maturation. Frontiers in Endocrinology 4: 2013.

65. Borgonovo B, Ouwendijk J, and Solimena M. Biogenesis of secretory granules. Current opinion in cell biology 18: 365–370, 2006.

66. Luo HR, Saiardi A, Nagata E, Ye K, Yu H, Jung TS, Luo X, Jain S, Sawa A, and Snyder SH. GRAB: a physiologic guanine nucleotide exchange factor for Rab3A, which interacts with inositol hexakisphosphate kinase. Neuron 31: 439–451, 2001.

67. Li H, Datunashvili M, Reyes RC, and Voglmaier SM. Inositol hexakisphosphate kinases differentially regulate trafficking of vesicular glutamate transporters 1 and 2. Front Cell Neurosci 16: 926794, 2022.

68. Štepihar D, Florke Gee RR, Hoyos Sanchez MC, and Fon Tacer K. Cell-specific secretory granule sorting mechanisms: the role of MAGEL2 and retromer in hypothalamic regulated secretion. Frontiers in Cell and Developmental Biology Volume 11–2023: 2023.

69. Cutler AA, Corbett AH, and Pavlath GK. Biochemical Isolation of Myonuclei from Mouse Skeletal Muscle Tissue. Bio Protoc 7: 2017.

70. Adamovich Y, Ezagouri S, Dandavate V, and Asher G. Monitoring daytime differences in moderate intensity exercise capacity using treadmill test and muscle dissection. STAR Protocols 2: 100331, 2021.

